# Retrograde axonal autophagy and endocytic pathways are parallel but separate in neurons

**DOI:** 10.1101/2022.07.01.498478

**Authors:** Vineet Vinay Kulkarni, Max Henry Stempel, Anip Anand, David Kader Sidibe, Sandra Maday

**Affiliations:** Department of Neuroscience, Perelman School of Medicine at the University of Pennsylvania, Philadelphia, PA, 19104, USA

**Keywords:** Autophagy, axon, endosome, LC3, lysosome, neurons, α-synuclein

## Abstract

Autophagy and endocytic trafficking are two key pathways that regulate the composition and integrity of the neuronal proteome. Alterations in these pathways are sufficient to cause neurodevelopmental and neurodegenerative disorders. Thus, defining how autophagy and endocytic pathways are organized in neurons remains a key area of investigation. These pathways share many features and converge on lysosomes for cargo degradation, but what remains unclear is the degree to which the identity of each pathway is preserved in each compartment of the neuron. Here, we elucidate the degree of intersection between autophagic and endocytic pathways in axons of primary neurons. Using microfluidic chambers, we labeled newly-generated bulk endosomes and signaling endosomes in the distal axon, and systematically tracked their trajectories, molecular composition, and functional characteristics relative to autophagosomes. We find that newly-formed endosomes and autophagosomes both undergo retrograde transport in the axon, but as distinct organelle populations. Moreover, these pathways differ in their degree of acidification and association with molecular determinants of organelle maturation. These results suggest that the identity of autophagic and newly endocytosed organelles is preserved for the length of the axon. Lastly, we find that expression of a pathogenic form of α-synuclein, a protein enriched in presynaptic terminals, increases merging between autophagic and endocytic pathways. Thus, aberrant merging of these pathways may represent a mechanism contributing to neuronal dysfunction in Parkinson’s disease and related α-synucleinopathies.

**SIGNIFICANCE STATEMENT:** Autophagy and endocytic trafficking are retrograde pathways in neuronal axons that fulfill critical degradative and signaling functions. These pathways share many features and converge on lysosomes for cargo degradation, but the extent to which the identity of each pathway is preserved in axons is unclear. We find that autophagosomes and endosomes formed in the distal axon undergo retrograde transport to the soma in parallel but separate pathways. These pathways also have distinct maturation profiles along the mid-axon, further highlighting differences in the potential fate of transported cargo. Strikingly, expression of a pathogenic variant of α-synuclein increases merging between autophagic and endocytic pathways, suggesting that mis-sorting of axonal cargo may contribute to neuronal dysfunction in Parkinson’s disease and related α-synucleinopathies.

## INTRODUCTION

Autophagy and endocytic trafficking are two key pathways that regulate the composition and integrity of the neuronal proteome (Winckler et al., 2018; Sidibe et al., 2022). Dysfunction in autophagic and endolysosomal trafficking is linked to many neurodevelopmental and neurodegenerative diseases (Winckler et al., 2018; Malik et al., 2019; Sidibe et al., 2022). But how autophagy and endocytic pathways are organized in the context of neurons to maintain protein and organelle homeostasis is a major area of investigation. Live-cell imaging of these organelle populations in axons has revealed that these pathways have many features in common.

Autophagosomes and endosomes are both formed in the distal axon at presynaptic terminals (Hollenbeck, 1993; Overly and Hollenbeck, 1996; Maday et al., 2012; Maday and Holzbaur, 2014; Soukup et al., 2016; Stavoe et al., 2016; Hill et al., 2019). Following formation, these organelles undergo retrograde transport and deliver cargos to the soma for degradation by resident lysosomes (Hollenbeck, 1993; Overly and Hollenbeck, 1996; Lee et al., 2011; Maday et al., 2012; Maday and Holzbaur, 2014). Thus, autophagic and endocytic pathways have similar trafficking trajectories in axons. However, the degree to which these pathways merge and at what point they merge is poorly understood. Endosomes transport a variety of cargos, some of which are destined for degradation whereas others need to be preserved for functional purposes (e.g. signaling). Thus, what is the trafficking itinerary of endocytosed cargos? Do endocytic cargos merge with autophagosomes for degradation?

Several studies established that axonal autophagosomes share many molecular signatures of endosomal organelles, suggesting considerable overlap between autophagic and endocytic pathways in axons. After autophagosomes form in axon terminals, they become positive for the Rab7 GTPase and the lysosomal-associated membrane protein 1 (LAMP1), components of late endosomes and lysosomes (Lee et al., 2011; Maday et al., 2012; Cheng et al., 2015; Maday and Holzbaur, 2016). Moreover, autophagosomes become partially acidified (Lee et al., 2011; Maday et al., 2012). Thus, autophagosomes formed in the distal axon are proposed to mature into amphisomes (products of fusion between autophagosomes and endosomal organelles), and this maturation may trigger their retrograde transport to the soma (Cheng et al., 2015; Farias et al., 2017; Farfel-Becker et al., 2019; Lie et al., 2021). Some studies suggest that a population of axonal autophagosomes fuses with signaling endosomes, a subset of endosomes that carry neurotrophin-mediated signaling information in the form of active BDNF-TrkB complexes (Kononenko et al., 2017; Andres-Alonso et al., 2019). Retrograde transport of these “signaling amphisomes” may be important for neuronal development and function (Kononenko et al., 2017; Andres-Alonso et al., 2019). Lastly, foundational studies from Hollenbeck and colleagues reported that long-term incubation with fluid-phase endocytic substrates results in their accumulation in axonal autophagosomes (Hollenbeck, 1993). But the degree of overlap between autophagosomes and newly-endocytosed cargo in the axon remains unclear.

Here, we set out to determine the extent of overlap between autophagic and endocytic pathways in axons of primary neurons. We cultured primary neurons in compartmentalized microfluidic chambers to selectively label only newly-formed endosomes in the distal axon. We applied different cargo molecules to label a broad population of endosomes that would include cargos to be degraded, and a subpopulation of signaling endosomes that carry cargos to be preserved (i.e. BDNF). Using live-cell imaging, we systematically tracked the trajectories, molecular identity, and functional characteristics of these organelles in the axon at high temporal resolution. We find that autophagosomes and newly-formed endosomes both undergo retrograde transport in the axon, but as distinct organelle populations. Moreover, these pathways differ in their extent of acidification and association with late endosomal markers, suggesting distinct routes of organelle maturation. In fact, newly endocytosed cargos in the axon do not reach degradative compartments until the soma. Lastly, we find that expression of a pathogenic form of α-synuclein, a protein enriched in presynaptic terminals, increases merging between autophagic and endocytic pathways. Combined, these results suggest that the identity of autophagic and newly endocytosed organelles is preserved for the length of the axon. Moreover, disruption of these pathways may lead to missorting of endocytic cargos and represents a possible mechanism contributing to neuronal dysfunction in Parkinson’s disease and related α-synucleinopathies.

## MATERIALS AND METHODS

### Reagents and materials

Transgenic mice expressing GFP-LC3 (EGFP fused to the C terminus of rat LC3B) were obtained from the RIKEN BioResource Research Center (RBRC00806; strain B6.Cg-Tg (CAG-EGFP/LC3)53Nmi/NmiRbrc; GFP-LC3#53) and maintained as heterozygotes. All animal protocols were approved by the Institutional Animal Care and Use Committee at the University of Pennsylvania. Constructs included GFP-Rab7 (Addgene #12605), LAMP1-RFP (Addgene #1817), EGFP-α-synuclein-WT (Addgene #40822), and EGFP-α-synuclein-A53T (Addgene #40823). BSA conjugates included BSA-488 (Thermo Fisher Scientific/Molecular Probes; A13100), BSA-647 (Thermo Fisher Scientific/Molecular Probes; A34785), DQ-Green-BSA (Thermo Fisher Scientific/Molecular Probes; D12050), and DQ-Red-BSA (Thermo Fisher Scientific/Molecular Probes; D12051). To label acidic compartments, Lysotracker Green (Thermo Fisher Scientific; L7526) and Lysotracker Deep-Red dyes (Thermo Fisher Scientific; L12492) were used. For labeling signaling endosomes, human BDNF-biotin (Alomone labs; B-250-B) was conjugated to Quantum dots-605 (QD-605) (Thermo Fisher Scientific; Q10103MP). Primary antibodies for immunofluorescence included rabbit anti-glial fibrillary acidic protein (GFAP) (EMD Millipore; AB5804), rabbit anti-microtubule associated protein-2 (MAP2) (Millipore Sigma AB5622), and mouse anti-TUBB3/β3 tubulin (Novus Biologicals; MAB1195-SP). Secondary antibodies for immunofluorescence included goat anti-rabbit Alexa Fluor 488 (Thermo Fisher Scientific/Invitrogen; A11034) and goat anti-mouse Alexa Fluor 647 (Thermo Fisher Scientific/Invitrogen; A32728). Hoechst 33342 (Molecular probes; H3570) was used to label nuclei. Microfluidic chambers with 900 µm long microgrooves (XONA Microfluidics; RD900) were used in all experiments.

### Neuron-astrocyte co-culture in microfluidic chambers

Prior to neuronal dissection, 35mm glass-bottom fluorodishes (World Precision Instruments; FD35-100) were coated with 1 mg/ml poly-L-lysine (Peptide International; OKK-3056) in borate buffer for overnight in a 37°C, 5% CO_2_ incubator, and washed twice with TC-grade water (Fisher Lonza; BW17724Q) the next day. Water was carefully aspirated from the fluorodishes to make them completely dry. Sterilized and scompletely dry XONA microfluidic chambers (RD900) were then carefully attached to the glass bottom of fluorodishes, as per manufacturer’s instructions. One side of the microfluidic chambers was labelled “proximal” (in which cells were plated) and the opposite side of the chambers was labelled “distal”, into which some axons would eventually extend in a stochastic manner (see Fig. 1A). Maintenance medium (Neurobasal medium [Thermo Fisher Scientific/Gibco; 21103–049] supplemented with 2% B-27 [Thermo Fisher Scientific/Gibco; 17504–044], 37.5 mM NaCl, 33 mM glucose [Sigma; G8769], 2 mM glutaMAX, and 100 U/ml penicillin and 100 µg/ml streptomycin) was added to only the proximal side first, to establish fluid flow across microgrooves. Fluorodishes containing microfluidic chambers and medium on proximal side were kept in the 37°C, 5% CO_2_ incubator overnight to ensure fluid flow through the microgrooves. Before plating neurons, maintenance medium was gently aspirated from the proximal and distal chambers, leaving medium present in the microgrooves.

**Figure 1.**
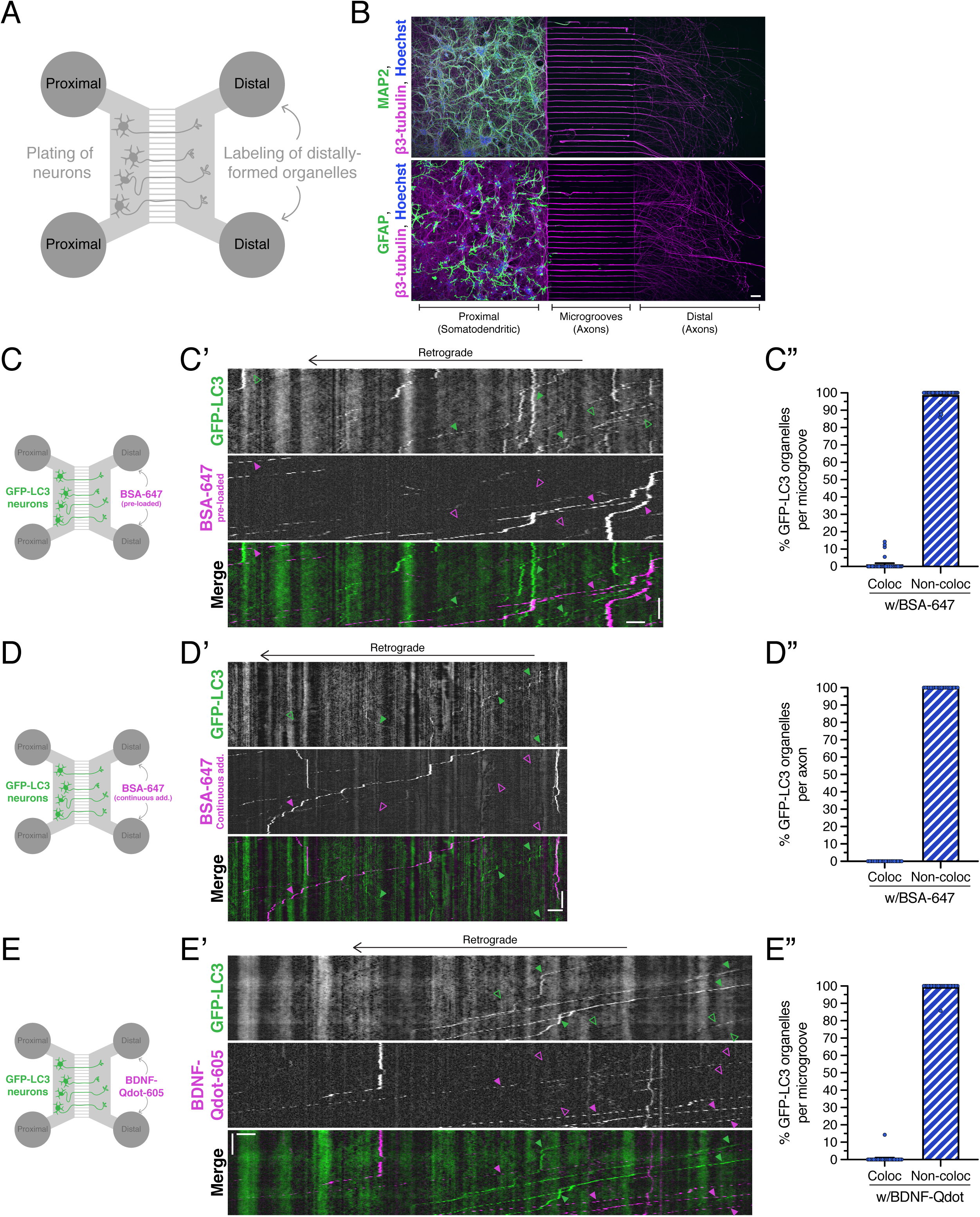
Autophagosomes and newly endocytosed cargo undergo retrograde transport as distinct organelle populations in the axon. **(A)** Schematic of microfluidic chamber. Neurons are plated in the proximal side of the chamber. Only axons can pass through narrow microgrooves to reach the distal chamber. Endocytic organelles forming in the distal axon can be selectively labeled by adding fluorescently-labeled cargos (e.g. BSA-647 or BDNF-Qdots) to only the distal chamber. Imaging is performed in the middle of the microgrooves. **(B)** Immunostain of dendrites (MAP2), neurons (β3-tubulin), astrocytes (GFAP), and total cells (Hoechst-positive nuclei) cultured in the microfluidic chamber. Bar, 100 µm. Neuronal somatodendritic compartments and astrocytes are restricted to the proximal chamber. Distal chamber contains only axons. **(C-C”)** Live cell imaging analysis of GFP-LC3-positive autophagosomes and newly endocytosed cargo labeled with BSA-647 in axons of primary cortical neurons. BSA-647 was added to only the distal chamber, and washed out prior to imaging in the mid-axon. **(C)** Schematic of experimental setup**. (C’)** Kymograph analysis of GFP-LC3 and BSA-647 motility in the axon. Vertical bar, 30 sec. Horizontal bar, 5 µm. Throughout the figure, solid arrowheads indicate organelles positive for the respective marker. Open arrowheads indicate organelles negative for the respective marker. Only GFP-LC3 transgenic neurons that have evidence of endocytosed BSA-647 are marked. **(C”)** Corresponding quantification of the percentage of autophagosomes positive or negative for BSA-647 in the axon (means ± SEM; N=35 microgrooves from 3 independent experiments; 8-9 DIV). **(D-D”)**. Live cell imaging analysis of GFP-LC3-positive autophagosomes and newly endocytosed cargo labeled with BSA-647 in axons of primary cortical neurons. BSA-647 was added to only the distal chamber and remained in the distal chamber during imaging to allow for continuous endocytic uptake. **(D)** Schematic of experimental setup**. (D’)** Kymograph analysis of GFP-LC3 and BSA-647 motility in the axon. Vertical bar, 1 min. Horizontal bar, 10 µm. Throughout the figure, solid arrowheads indicate organelles positive for the respective marker. Open arrowheads indicate organelles negative for the respective marker. Only GFP-LC3 transgenic neurons that have evidence of endocytosed BSA-647 are marked. **(D”)** Corresponding quantification of the percentage of autophagosomes positive or negative for BSA-647 in the axon (means ± SEM; N=20 axons from 4 independent experiments; 8-9 DIV). **(E-E”)** Live cell imaging analysis of GFP-LC3-positive autophagosomes and endocytosed BDNF-labeled Qdots-605 in axons of primary cortical neurons. BDNF-Qdots were added to only the distal chamber. **(E)** Schematic of experimental setup**. (E’)** Kymograph analysis of GFP-LC3 and BDNF-Qdot motility in the axon. Vertical bar, 30 sec. Horizontal bar, 5 µm. Throughout the figure, solid arrowheads indicate organelles positive for the respective marker. Open arrowheads indicate organelles negative for the respective marker. Only GFP-LC3 transgenic neurons that have evidence of endocytosed BDNF-Qdots are marked. **(E”)** Corresponding quantification of the percentage of autophagosomes positive or negative for BDNF-Qdots in the axon (means ± SEM; N=25 microgrooves from 3 independent experiments; 9-11 DIV).

For neuronal culture, cerebral cortices were dissected from brains of GFP-LC3 transgenic mouse embryos, or non-transgenic wild type (WT) mouse embryos of either sex at day 15.5; detailed methods are found in Dong et al., 2019 (Dong et al., 2019). In brief, tissue was digested with 0.25% trypsin for 10 min at 37°C, and then triturated through a small-bore glass Pasteur pipette to achieve a homogeneous cell suspension. For experiments requiring WT neurons (as in Fig. 1B, Fig. 2B-C, Fig. 3-5, Extended Data Fig. 2-1), 100,000 WT neurons were plated in a low volume (<30 µl) of attachment medium (Minimal Essential Media supplemented with 10% heat inactivated horse serum, 33 mM glucose, 1 mM pyruvic acid, and 37.5 mM NaCl) in the top well of the proximal side of the microfluidic chambers. Plating cells in low volume (<30 µl) is essential to retain most cells in the channel connecting the two wells on the proximal side, and thus increasing the probability of neurons sending axons through the microgrooves to the distal side. For experiments requiring GFP-LC3 transgenic neurons, GFP-LC3 neurons were diluted with WT neurons to achieve single axon resolution. For Fig. 1C, Fig. 1E, and Fig. 2A, 25,000 GFP-LC3 transgenic neurons were diluted with 75,000 WT neurons (totalling 100,000 neurons) and plated in the top well of the proximal side of the chamber. For Fig. 1D, 5,000 GFP-LC3 transgenic neurons were diluted with 95,000 WT neurons (totaling 100,000 neurons) and plated in the top well of the proximal side of the microfluidic chamber. Neurons were allowed to attach to the coverslip for ∼30 minutes, and then 100 µl of maintenance medium each was added to each of the 4 wells of the microfluidic chamber. Cells were maintained in this medium until they were DIV2. On DIV2, neurons were co-cultured with astrocytes, as detailed below.

**Figure 2.**
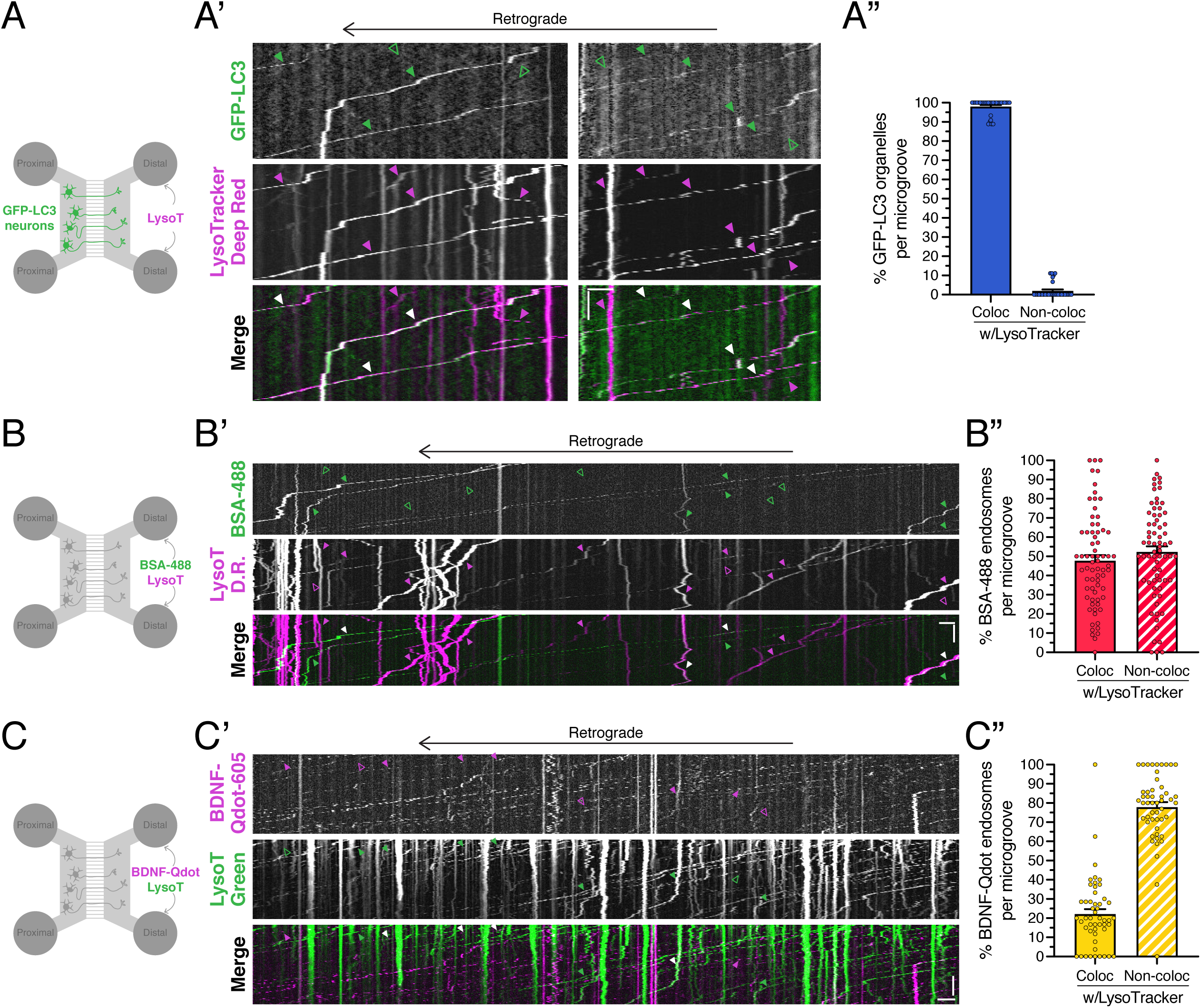
Autophagosomes and organelles containing newly endocytosed cargo exhibit distinct maturation rates. **(A-A”)** Live cell imaging analysis of GFP-LC3-positive autophagosomes and Lysotracker-Deep Red in axons of primary cortical neurons. Lysotracker-Deep Red was added to only the distal chamber. **(A)** Schematic of experimental setup. (**A’**) Kymograph analysis of GFP-LC3 and Lysotracker-Deep Red motility in the axon. Vertical bar, 30 sec. Horizontal bar, 5 µm. Throughout the figure, solid arrowheads indicate organelles positive for the respective marker. Open arrowheads indicate organelles negative for the respective marker. Solid white arrowheads in the merged image indicate organelles co-positive for both markers. **(A”)** Corresponding quantification of the percentage of autophagosomes positive or negative for Lysotracker-Deep Red in the axon (means ± SEM; N=30 microgrooves from 3 independent experiments; 9-10 DIV). **(B-B”)** Live cell imaging analysis of newly endocytosed BSA-488 and Lysotracker-Deep Red in axons of primary cortical neurons. BSA-488 and Lysotracker-Deep Red were applied to only the distal chamber. **(B)** Schematic of experimental setup. **(B’)** Kymograph analysis of BSA-488 and Lysotracker-Deep Red motility in the axon. Vertical bar, 30 sec. Horizontal bar, 5 µm. **(B”)** Corresponding quantification of the percentage of BSA-488 puncta positive or negative for Lysotracker-Deep Red in the axon (means ± SEM; N=70 microgrooves from 4 independent experiments; 8-9 DIV). **(C-C”)** Live cell imaging analysis of newly endocytosed BDNF-Qdots-605 and Lysotracker-Green in axons of primary cortical neurons. BDNF-Qdots-605 and Lysotracker-Green were added to only the distal chamber. **(C)** Schematic of experimental setup. **(C’)** Kymograph analysis of BDNF-Qdot-605 and Lysotracker-Green motility in the axon. Vertical bar, 30 sec. Horizontal bar, 5 µm. The three white arrowheads in the top left of the merged kymograph denote three different regions of the same track. **(C”)** Corresponding quantification of the percentage of BDNF-Qdot puncta positive or negative for Lysotracker-Green in the axon (means ± SEM; N= 50 microgrooves from 3 independent experiments; 8-9 DIV).

**Figure 3.**
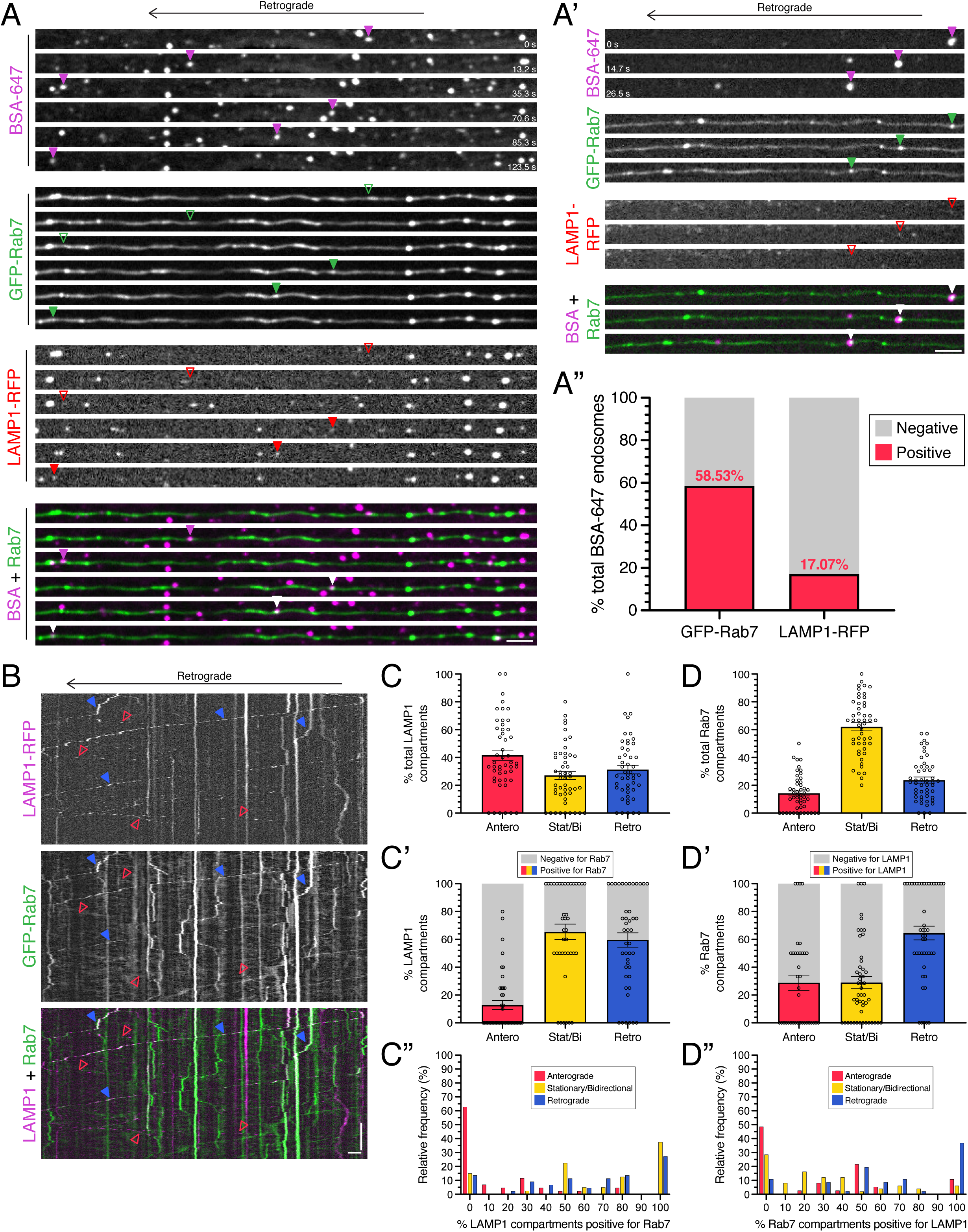
Newly-formed endosomes have distinct molecular signatures of maturation. **(A-A”).** Live cell imaging analysis of GFP-Rab7, LAMP1-RFP, and newly endocytosed BSA-647 in the axon of primary cortical neurons. BSA-647 was added to only the distal chamber, and present during the imaging. **(A)** Time series of kymostacks depicting an example of a BSA-positive organelle that is negative for Rab7 and LAMP1, and an example of a BSA-positive organelle co-positive for Rab7 and LAMP1. Throughout the figure, solid arrowheads indicate organelles positive for the respective marker. Open arrowheads indicate organelles negative for the respective marker. Solid white arrowheads in the merged image indicate organelles co-positive for both markers. Bar, 5 µm. **(A’)** Time series of kymostacks depicting an example of a BSA-positive organelle that is positive for Rab7, but negative for LAMP1. Bar, 5 µm. **(A”)** Quantitation of the percentage of newly formed endosomes (labeled with BSA-647) that are co-positive for Rab7 or LAMP1 (percentage based on a pool of 41 BSA-647 puncta counted from 22 axons from 5 independent experiments; for the 7 BSA-647 puncta that are LAMP1-positive, 5 are Rab7-positive and 2 are Rab7-negative). **(B)** Kymograph of LAMP1-RFP and GFP-Rab7 motility in the axon of primary cortical neurons. Solid blue arrowheads depict retrograde LAMP1 compartments that are co-positive for Rab7. Open red arrowheads depict anterograde LAMP1 compartments that are negative for Rab7. Vertical bar, 1 min. Horizontal bar, 5 µm. **(C)** Quantification of the percentage of LAMP1 compartments moving either anterograde, retrograde, or stationary/bidirectional (means ± SEM; N=49 axons from 5 independent experiments; 9 DIV). **(C’-C”)** Quantification of the percentage of LAMP1 compartments moving either anterograde, retrograde, or stationary/bidirectional that are co-positive for Rab7 (C’, means ± SEM; C”, histogram; N=40-44 axons from 5 independent experiments; 9 DIV). **(D)** Quantification of the percentage of Rab7 compartments moving either anterograde, retrograde, or stationary/bidirectional (means ± SEM; N=49 axons from 5 independent experiments; 9 DIV). **(D’-D”)** Quantification of the percentage of Rab7 compartments moving either anterograde, retrograde, or stationary/bidirectional that are co-positive for LAMP1 (D’, means ± SEM; D”, histogram; N=37-49 axons from 5 independent experiments; 9 DIV).

**Figure 5.**
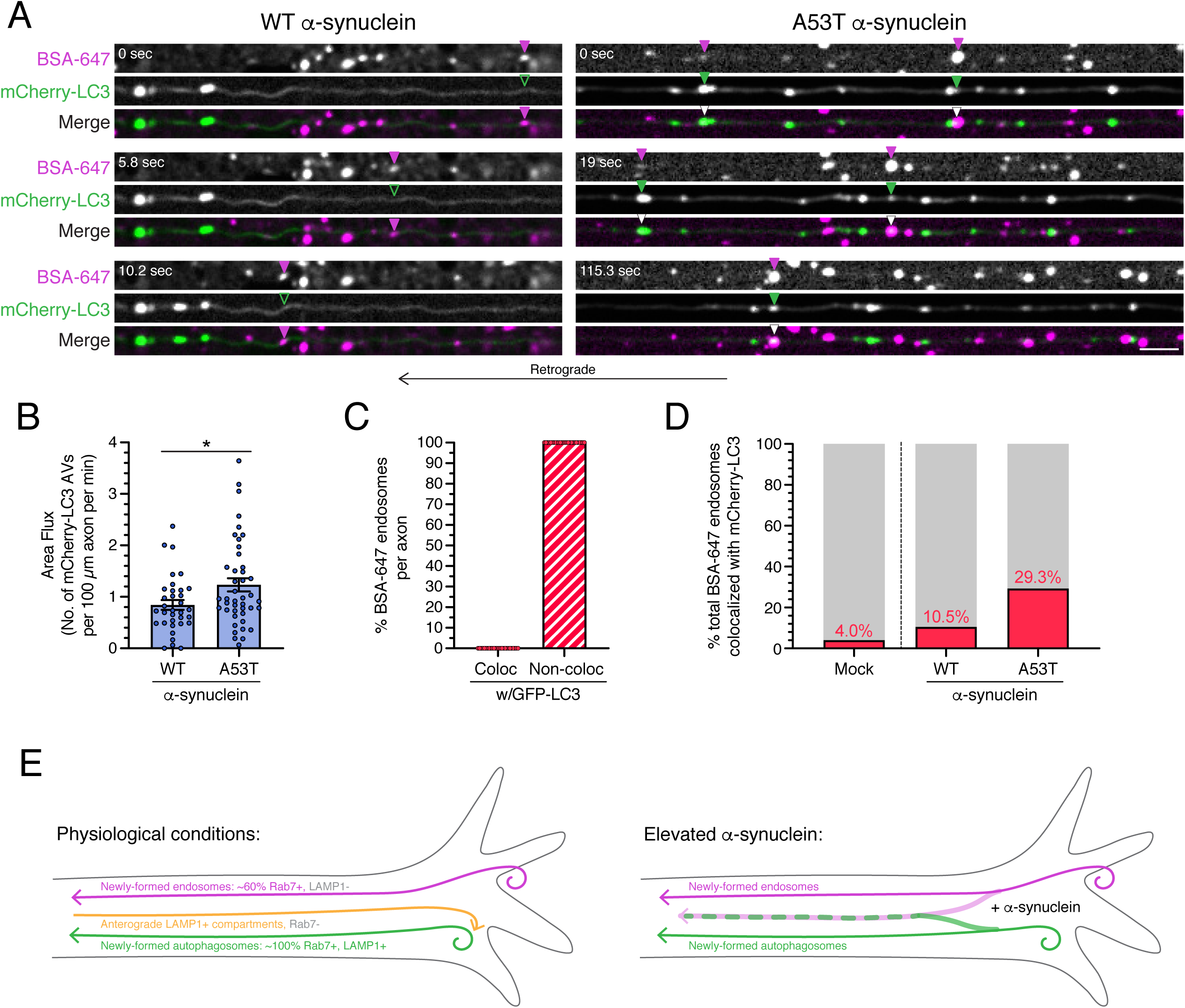
Expression of pathogenic α-synuclein increases merging of retrograde pathways for autophagy and endocytic trafficking. **(A-B, D)** Live cell imaging of newly endocytosed BSA-647 and mCherry-LC3 in axons of primary cortical neurons expressing GFP-tagged WT or A53T α-synuclein. BSA-647 was added to only the distal chamber, and present for the imaging. **(A)** Time series of kymostacks showing an increase in colocalization of newly endocytosed BSA-647 with mCherry-LC3 in the presence of A53T α-synuclein as compared with WT α-synuclein. Solid arrowheads indicate organelles positive for the respective marker. Open arrowheads indicate organelles negative for the respective marker. Solid white arrowheads in the merged image indicate organelles co-positive for both markers. Bar, 5 µm. **(B)** Corresponding quantitation of mCherry-LC3 area flux (means ± SEM; unpaired t-test; *, p ≤ 0.05; n=34-43 axons from 4 independent experiments; 9-12 DIV). **(C)** Live cell imaging analysis of GFP-LC3-positive autophagosomes and newly endocytosed cargo labeled with BSA-647 in axons of primary cortical neurons. BSA-647 was added to only the distal chamber and remained in the distal chamber for imaging to allow for continuous endocytic uptake. Quantitation of the percentage of newly formed endosomes as labeled with BSA-647 that are positive for GFP-LC3 in the axon (means ± SEM; n=26 axons from 4 independent experiments; 8-9 DIV). Dataset is the same as the dataset quantitated in Figure 1D-D”, but the data are quantified from the BSA perspective. **(D)** Corresponding quantitation of the experiment in **(A)** of the percentage of BSA-positive puncta co-positive for Cherry-LC3 in axons transfected with mock conditions, or plasmids encoding WT or A53T α-synuclein (mock, percentage based on a pool of 50 BSA puncta counted from 22 axons from 4 independent experiments; 10-11 DIV; WT α-synuclein, percentage based on a pool of 38 BSA puncta counted from 13 axons from 4 independent experiments; 9-12 DIV; A53T α-synuclein, percentage based on a pool of 58 BSA puncta counted from 24 axons from 4 independent experiments; 9-12 DIV). **(E)** Model schematics depicting the parallel pathways for autophagy and endocytic trafficking in the distal axon under physiological conditions and in elevated α-synuclein. Prior studies have established the extensive overlap between autophagosomes and Rab7-LAMP (Lee et al., 2011; Maday et al., 2012; Cheng et al., 2015; Kulkarni et al., 2021).

Astrocytes were prepared by dissecting cerebral cortices from brains of non-transgenic neonatal mice of either sex at P0-P1. The meninges were removed and the tissue was digested with 0.25% trypsin (Thermo Fisher Scientific/Gibco; 15090–046) for 10 min at 37°C, then triturated with a 5 ml pipette to break up large pieces of tissue, and then triturated with a P1000 pipet until a homogeneous cell suspension is achieved. Cells were passed through a strainer with 40 µm pores (Falcon; 352340) and plated at a density of 2,000,000–3,000,000 astrocytes per 10 cm dish. Astrocytes were grown in glial media (DMEM [Thermo Fisher Scientific/Gibco; 11965–084] supplemented with 10% heat inactivated fetal bovine serum [Hyclone; SH30071.03], 2 mM Glutamax [Thermo Fisher Scientific/Gibco; 35050–061], 100 U/ml penicillin and 100 µg/ml streptomycin [Thermo Fisher Scientific/Gibco; 15140–122]) at 37°C in a 5% CO_2_ incubator. The next day after plating, and every 3–4 d following, astrocytes were fed by replacing glial media. When astrocytes reached 80–90% confluence (∼8–10 d after plating), astrocytes were trypsinized and used for co-culturing experiments. After dissociation from the 10-cm plate, trypsin was inactivated using glial media. In some cases, astrocytes were passaged once before co-culturing with neurons. 10,000 astrocytes (per 100,000 neurons) were then resuspended in co-culture media (Neurobasal supplemented with 2% B-27, 1% G-5, 0.25% GlutaMAX, 100 U/ml penicillin, and 100 µg/ml streptomycin) and plated on the neuron culture in the top well of the proximal side of the microfluidic chamber, and maintained at 37°C in a 5% CO_2_ incubator. After 2 days of co-culture, medium was aspirated from both sides of the microfluidic chamber, and 100 µl of co-culture medium containing 2 µM AraC (antimitotic drug, C6645; Sigma-Aldrich) was added to each well of the microfluidic chamber. Addition of AraC was done to prevent cell division and promote maturation of the astrocytes. The co-culture was fed every other day by removing 50 µl of medium from each well of the 4 wells, and adding 80 µl of fresh co-culture medium containing 2 µM AraC to each of the 4 wells. For experiments in Fig. 3, neuron-astrocyte co-cultures were transfected in the proximal chamber with both GFP-Rab7 and LAMP1-RFP using Lipofectamine 2000 after 4 days of co-culture (DIV6 total for neurons), as per manufacturer’s instructions. In the same way, for experiments in Fig. 5, neurons were transfected with either EGFP-α-synuclein-WT or EGFP-α-synuclein-A53T.

### Live-cell confocal imaging

Live-cell imaging of organelle dynamics was performed on a BioVision spinning disk confocal microscope system consisting of a Leica DMi8 inverted widefield microscope, a Yokagawa W1 spinning disk confocal microscope, and a Photometrics Prime 95B scientific complementary metal–oxide–semiconductor camera. Images were acquired with VisiView software using a 63X/1.4 NA Plan Apochromat oil-immersion objective and solid-state 405-, 488-, 561-, and 640-nm lasers for excitation. The microscope was equipped with an environmental chamber at 37°C and adaptive focus control to maintain a constant focal plane during live-cell imaging. Each microfluidic chamber was imaged for a maximum of 30 minutes on the scope.

### Immunostaining of co-cultured neurons and astrocytes in microfluidic chambers

On DIV8 of neuronal culture (DIV6 of the co-culture), microfluidic chambers were carefully removed from the fluorodishes using forceps without disrupting the underlying layer of cells and axons in microgrooves. Samples were fixed for 10 min in 4% PFA/4% sucrose in PBS (150 mM NaCl, 50 mM NaPO_4_, pH 7.4) previously warmed to 37°C. Cells were washed two times in PBS, permeabilized for 5 min in 0.1% Triton X-100 (Thermo Fisher Scientific; BP151-100) in PBS, washed two times in PBS, and then blocked for 1 hr in PBS supplemented with 5% goat serum (Sigma-Aldrich; G9023) and 1% BSA (Thermo Fisher Scientific; BP1605-100). Cells were then incubated in primary antibody diluted in block solution for 1 hr at room temperature, washed three times for 5 min each in PBS, and then incubated in secondary antibody and Hoechst 33342 (Thermo Fisher Scientific /Molecular Probes; H3570) diluted in block solution for 1 hr at room temperature. Following incubation in secondary antibody, samples were washed three times for 5 min each in PBS. Samples were kept in the 3^rd^ PBS wash and imaged with the BioVision spinning-disk confocal microscope system described above using a 10X/0.32 NA Plan Fluotar dry objective and 405, 488- and 640nm lasers for excitation. Tiled Z-stacks were obtained that spanned the entire length (in the X direction) of the field of neurons and astrocytes in the microfluidic chambers; sections were taken every 1 µm in the Z-direction that spanned the entire depth of cells. Maximum projections of each Z-stack were made in FIJI using the maximum intensity for each pixel. Maximum projections from all three channels were merged to form a composite. Max projection composites for the proximal, microgroove, and distal sections of the microfluidic chamber were then stitched together using the FIJI “pairwise stitching” plugin.

### Fluidic isolation of dyes and reagents

Microfluidic chambers allow for fluidic isolation of dyes and reagents to one specific compartment (proximal or distal) by generating a hydrostatic pressure differential between the two compartments. To maintain fluidic isolation of a dye or reagent on one side, a pressure differential of 60-80 µl of medium was maintained between the two compartments. For example, to maintain fluidic isolation of a dye or reagent on the distal side, a total of 140 µl of medium containing the appropriate concentration of dye or reagent was added to the distal side and a total of 220 µl medium devoid of the dye or reagent was added to the proximal side (pressure differential of 220 µl – 140 µl = 80 µl more volume on the proximal versus distal side).

#### Fluidic isolation of BSA and Lysotracker Dyes

For Fig. 1C-D, Fig. 2B, Fig. 3A, and Fig. 5A, at DIV 8-10 of neuronal culture, (DIV 9-12 for Fig. 5A), maintenance medium containing 24.4 mg/ml BSA-647 (or BSA-488 for Fig. 2B) was added to only the distal side of the microfluidic chamber while maintaining fluidic isolation, for 2 hours. For Fig. 2B, for the final 30 minutes of the 2-hour incubation, medium from the distal side was replaced with maintenance medium containing 100 nM Lysotracker Deep-Red along with 24.4 mg/ml BSA-488 while maintaining fluidic isolation. After the 2-hour incubation, for Fig. 1C and Fig. 2B, both proximal and distal chambers were washed with HibE imaging medium (Hibernate E [BrainBits] supplemented with 2% B-27, 2 mM GlutaMAX, 100 U/ml penicillin and 100 µg/ml streptomycin [Thermo Fisher Scientific/Gibco; 15140–122]) twice while maintaining fluidic isolation. Fresh HibE imaging medium was then added to both proximal and distal sides while maintaining fluidic isolation. For Fig. 1D, Fig. 3A, and Fig. 5A, medium on the distal side was replaced with HibE imaging medium containing 24.4 mg/ml BSA-647, and maintenance medium on the proximal side was replaced with HibE imaging medium, while maintaining fluidic isolation. We then imaged axons in the middle of the microgrooves of the microfluidic chambers, ∼100 µm away from the proximal and distal ends of the microgrooves, that displayed clear and distinct puncta (for transgenic or transfected markers or dyes). Imaging was performed at 1 fps for 2 minutes (Fig. 1C, Fig. 2B), at 1 fps for 5 minutes (Fig. 1D, and the mock transfected control quantified in Fig. 5D), at 0.68-0.89 fps for 5.61-7.36 minutes (Figs 3A), or at 0.76-0.89 fps for 6.61-7.36 minutes (Fig. 5A).

For Fig. 2A, at DIV 9-10 of neuronal culture, maintenance medium containing 100 nM Lysotracker Deep Red was added to the distal side for 30 minutes while maintaining fluidic isolation. After incubation, both proximal and distal chambers were washed with HibE imaging medium twice while maintaining fluidic isolation. Fresh HibE imaging medium was then added to both proximal and distal chambers while maintaining fluidic isolation. We then imaged axons in the mid-grooves of the microfluidic chambers, ∼100 µm away from the proximal and distal end of the microgrooves, that displayed clear and distinct puncta. Images were acquired at 1 fps for 2 minutes.

For Fig. 4A-B, at DIV 8-9 of neuronal culture, maintenance medium containing 24.4 mg/ml BSA-647 and 24.4 mg/ml DQ-Green-BSA was added to the distal side of the microfluidic chamber, and maintenance medium containing 24.4 mg/ml DQ-Green-BSA was added to the proximal side of the microfluidic chamber while maintaining fluidic isolation for 2 h (Fig. 4A) or 4 h (Fig. 4B). After incubation, both proximal and distal chambers were washed with HibE imaging medium twice while maintaining fluidic isolation. Fresh HibE imaging medium was then added to both proximal and distal sides while maintaining fluidic isolation. We imaged axons in the mid-grooves of the microfluidic chambers, ∼100 µm away from the proximal and distal ends of the microgrooves, that displayed clear and distinct puncta in either channel. Images were acquired at 1 fps for 1 minute. On the proximal side, cell somas containing clear and distinct puncta were imaged at 1 fps for 1 minute.

**Figure 4.**
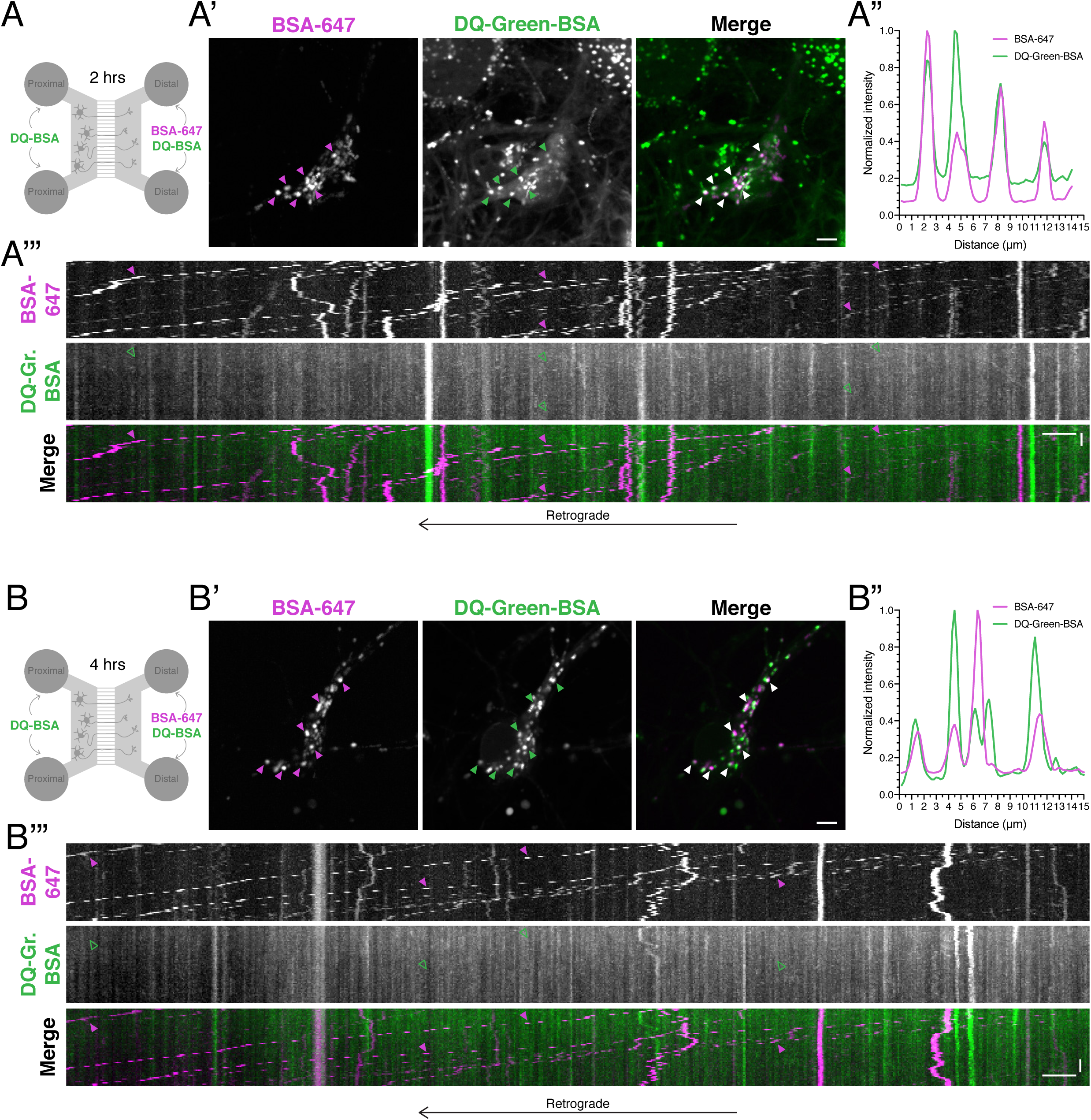
Newly endocytosed cargos in the axon are not in a degradative compartment until reaching the soma. **(A-A’”)** Live cell imaging analysis of newly endocytosed BSA-647 and DQ-Green-BSA in the soma and axon of primary cortical neurons after 2 hr of incubation with BSA conjugates (8-9 DIV). DQ-Green-BSA was added to both proximal and distal chambers, but BSA-647 was added to only the distal chamber. **(A)** Schematic of experimental setup. **(A’-A”)** *En face* images of the soma and corresponding line scans of BSA-647 and DQ-Green-BSA in the soma. Bar, 5 µm. Throughout the figure, solid arrowheads indicate organelles positive for the respective marker. Open arrowheads indicate organelles negative for the respective marker. Solid white arrowheads in the merged image indicate organelles co-positive for both markers. **(A’”)** Kymograph analysis of newly endocytosed BSA-647 and DQ-Green-BSA motility in the axon. Vertical bar, 12 sec. Horizontal bar, 5 µm. **(B-B’”)** Live cell imaging analysis of newly endocytosed BSA-647 and DQ-Green-BSA in the soma and axon of primary cortical neurons after 4 hr of incubation with BSA conjugates (8-9 DIV). DQ-Green-BSA was added to both proximal and distal chambers, but BSA-647 was added to only the distal chamber. **(B)** Schematic of experimental setup. **(B’-B”)** *En face* images of the soma and corresponding line scans of BSA-647 and DQ-Green-BSA in the soma. Bar, 5 µm. **(B’”)** Kymograph analysis of newly endocytosed BSA-647 and DQ-Green-BSA motility in the axon. Vertical bar, 12 sec. Horizontal bar, 5 µm.

For Extended Data Fig. 2-1A, at DIV 8-11 of neuronal culture, maintenance medium containing 24.4 mg/ml DQ-Green-BSA and DQ-Red-BSA was added only to the proximal side of the microfluidic chamber for 2 or 4 hours, while maintaining fluidic isolation. After incubation, both proximal and distal chambers were washed with HibE imaging medium twice while maintaining fluidic isolation. Fresh HibE imaging medium was then added to both proximal and distal sides while maintaining fluidic isolation. We then imaged axons in the mid-grooves of microfluidic chambers, ∼100 µm away from the proximal and distal ends, at 1 fps for 2 minutes. On the proximal side, cell somas containing clear and distinct puncta were imaged at 1 fps for 30 seconds.

For Extended Data Fig. 2-1B, at DIV 8-10 of neuronal culture, maintenance medium containing 24.4 mg/ml DQ-Red-BSA was added only to the proximal side of the microfluidic chamber for 4 hours, while maintaining fluidic isolation. For the final 30 minutes of the 4-hour incubation, medium from the proximal side was replaced with maintenance medium containing 100 nM Lysotracker Green along with 24.4 mg/ml DQ-Red-BSA while maintaining fluidic isolation. After incubation, both proximal and distal chambers were washed with HibE imaging medium twice while maintaining fluidic isolation. Fresh HibE imaging medium was then added to both proximal and distal sides while maintaining fluidic isolation. We then imaged axons in the mid-grooves of microfluidic chambers, ∼100 µm away from the proximal and distal ends, at 1 fps for 2 minutes. On the proximal side, cell somas containing clear and distinct puncta were imaged at 1 fps for 2 minutes.

For Extended Data Fig. 1-1, primary wild-type (WT) rat cortical neurons were obtained as a gift from Dr. Kelly Jordan-Sciutto (University of Pennsylvania, Philadelphia, PA), and isolated as previously described (Kulkarni et al., 2021). Microfluidic chambers were mounted on 25-mm acid-washed glass coverslips coated with 1 mg/ml poly-L-lysine (Peptide International; OKK-3056). 100,000 neurons were plated in rat neuron maintenance medium (Neurobasal medium supplemented with 2% B-27, 33 mM glucose, 100 U/ml penicillin, and 100 µg/ml streptomycin) in the top well of the proximal side of the microfluidic chambers. For these cultures, the anti-mitotic reagent AraC was not included in the culturing protocol. The culture was fed every day by removing 50 µl of medium and adding 65 µl of fresh medium from each well of the 4 wells. At DIV 8-14, maintenance medium containing 24.4 mg/ml BSA-647 was added to only the distal side (for Extended Data Fig. 1-1A) or only the proximal side (for Extended Data Fig. 1-1B) of the microfluidic chamber while maintaining fluidic isolation in the respective direction; neurons were incubated with BSA-647 for 2 hours. After incubation, both proximal and distal chambers were washed with HibE imaging medium twice while maintaining fluidic isolation in the respective direction. Fresh HibE imaging medium was then added to both proximal and distal sides while maintaining fluidic isolation in the respective direction. We then imaged axons in the mid-grooves of microfluidic chambers, ∼100 µm away from the proximal and distal ends, at 2 fps for 2-3 minutes.

#### Fluidic isolation of BDNF-Qdot-605

Human BDNF-biotin (50 nM) was incubated with 50 nM Qdot-605 Streptavidin conjugate (Thermo Fisher Scientific; Q10101MP) on ice for 1 h in Neurobasal medium to facilitate the biotin-streptavidin interaction, and generate BDNF-Qdot-605. For Fig. 1E and Fig. 2C, at DIV 9-11 of neuronal culture, maintenance medium containing 0.4 nM BDNF-Qdot-605 was added to only the distal side of the microfluidic chamber for 4 h while maintaining fluidic isolation. For Fig. 2C, for the final 30 minutes of BDNF-Qdot-605 treatment, fresh medium containing 0.4 nM BDNF-Qdot-605 and 100 nM Lysotracker Green was added to the distal side while maintaining fluidic isolation. After incubation, medium on both proximal and distal sides was replaced with HibE imaging medium while maintaining fluidic isolation. We then imaged axons in the mid-grooves of the microfluidic chambers, ∼100 µm away from the proximal and distal ends of the microgrooves. Images were acquired at 1 fps for 2 minutes.

### Image analysis

#### Tracking organelle dynamics and co-positivity in individual axons

Using the KymoToolBox plugin for Fiji (Hangen et al., 2018), kymographs were generated using line width 3 pixels for axons (Figs. 1C, 1D, 1E, and 2A). For co-transfected axons in Figs. 3A and 5A, the KymoToolBox plugin was used to generate straightened kymostacks of individual axons with a width of 20 pixels. In conditions of fluidic isolation maintained on the distal side, GFP-LC3 transgenic axons (or axons transfected with fluorescently-tagged constructs) present in microgrooves, that displayed at least one motile punctum labelled with the relevant dye, were considered for analysis. This criterion ensured that the GFP-LC3 transgenic or transfected axons had access to the dyes/reagents on the distal side.

For Figs. 1C, 1D, 1E and 2A, individual GFP-LC3-positive puncta were then determined to be co-positive for the respective marker if their signals overlapped and they co-migrated. Similarly, for Figs. 3A and 5A, individual BSA-647-positive puncta present in the axons of interest were determined to be co-positive for the respective fluorescent marker if their signals overlapped and they co-migrated. In all cases, both the kymograph as well as straightened kymostacks were considered while determining co-positivity. For Figs. 1C, 1E and 2A, due to the low number of puncta found in individual axons, GFP-LC3-positive puncta from all axons in a single microgroove that were co-positive for the respective marker were summed and plotted as a percentage of total GFP-LC3 organelles counted per microgroove. All GFP-LC3-positive puncta (for Fig. 1D) from single axons that were co-positive for the respective marker were summed and plotted as a percentage of total GFP-LC3 positive organelles counted per axon. Due to the low frequency of BSA-647-positive puncta traversing through transfected axons, the graphs in Fig. 3A and Fig. 5D were generated by determining the total number of BSA-647 puncta detected in transfected axons across all individual experiments that were co-positive for the respective marker of interest. Note that the “mock” control in Fig. 5D was performed independently from the WT-α-Synuclein and A53T-α-Synuclein conditions. The graph in Fig. 5C is generated using data from Fig. 1D-D’ by counting all BSA-647 puncta present in axons and determining the percentage of total BSA-647 puncta co-positive for GFP-LC3.

For Figs. 3B-D, kymographs were generated using a line width of 3 pixels using the KymoToolBox plugin, for axons co-transfected with LAMP1-RFP and GFP-Rab7. From each kymograph, along with consulting the movies, the percentage of LAMP1-RFP-positive puncta (Fig 3C) or GFP-Rab7-positive puncta (Fig 3D) moving in the net retrograde direction (≥1 µm displacement per 1 min) versus net anterograde direction (≥1 µm displacement per 1 min) was determined. Non-processive puncta that did not exhibit a displacement of 1 µm per 1 min were binned as showing bidirectional and stationary motility. For Fig. 3C’, within each category for direction, the LAMP1-RFP-positive puncta per axon were further categorized into puncta that were positive or negative for GFP-Rab7. Fig. 3C” displays a histogram representation of data in Fig. 3C’. Similarly, for Fig. 3D’, GFP-Rab7 positive puncta in each direction were further categorized into puncta that were positive or negative for LAMP1-RFP.

#### Tracking organelle dynamics and co-positivity in axon bundles in microgrooves

For experiments that did not use GFP-LC3 transgenic or transfected neurons, we were unable to discern single axon resolution. Thus, we quantified puncta detected within axon bundles present in the microgrooves. Using the KymoToolBox plugin, kymographs were generated in the microgrooves using a line width of 35 pixels (Figs. 2B, 2C, 4A, 4B, and Extended Data Fig. 2-1A-B) or a line width of 25 pixels (Extended Data Fig. 1-1). Microgrooves displaying clear puncta and motility of at least one marker were selected for analysis. For Extended Data Fig. 1-1, from each kymograph, along with consulting the movie, the percentage of BSA-647-positive puncta moving in the net retrograde direction (≥1 µm displacement per 1 min) versus net anterograde direction (≥1 µm displacement per 1 min) was determined. Non-processive puncta that did not exhibit a net displacement of 1 µm per 1 min were binned as showing bidirectional and stationary motility. For Figs. 2B and 2C, all unambiguous puncta positive for BSA-488 or BDNF-Qdot-605 in the microgroove were determined to be co-positive for Lysotracker Deep-Red or Lysotracker Green if their signals overlapped and they co-migrated. All BSA-488 or BDNF-QD-605 puncta in a microgroove that were co-positive for the relevant marker were then plotted as a percentage of total BSA-488 or BDNF-QD-605 endosomes counted per microgroove. For Extended Data Fig. 2-1B” and B””, only puncta that showed long-range motility in a retrograde or anterograde direction (1 µm displacement per 1 min) were considered for analysis. For Extended Data Fig. 2-1B”, the total number of DQ-Red-BSA that exhibited long-range movement in either anterograde or retrograde directions was pooled across all microgrooves in all experiments. Within each category for direction, the percentage of DQ-Red-BSA puncta that were co-positive for Lysotracker Green was then plotted out of total DQ-Red-BSA detected. The graph in Extended Data Fig. 2-1B”” was generated by determining the percentage of Lysotracker Green positive puncta undergoing long-range motility in either the anterograde or retrograde direction per microgroove.

#### Line scan analysis of neuronal somas

For graphs in Figs. 4A’’ and 4B’’, a segmented line of pixel width of 3 was drawn tracing BSA-647 positive puncta in the cell soma. This line was overlaid onto the DQ-Green-BSA channel. The gray value for each pixel along the line was obtained using the “Plot Profile” tool in FIJI. The raw gray values were then normalized to the maximum gray value within each respective channel along that line and plotted as a function of distance across the line.

#### Area flux analysis

For calculating the area flux of mCherry-LC3 positive AVs in Fig. 5B, kymographs of axons co-transfected with mCherry-LC3 and either EGFP-α-synuclein-WT or EGFP-α-synuclein-A53T were generated with a line-width of 3 pixels using the KymoToolBox plugin as described above. Consulting the kymographs as well as movies, the total number of mCherry-LC3 positive AVs appearing in each kymograph was determined, and normalized for both length of axon (per 100 µm) and time of movie (per min). The values generated were plotted as area flux.

#### Figure preparation and statistical analysis

All image measurements were obtained from the raw data. GraphPad Prism was used to plot graphs and perform statistical analyses; statistical tests are denoted within the figure legend. For presentation of images, maximum and minimum gray values were adjusted linearly in Fiji, and images were assembled in Adobe Illustrator.

## RESULTS

### Retrograde autophagosomes do not carry newly endocytosed cargos

Autophagosomes and endosomes regulate the axonal proteome by transporting cargos from the distal axon to the soma for degradation in lysosomes (Winckler et al., 2018; Sidibe et al., 2022). The degree of overlap between these pathways, however, is poorly understood. In this study, we use live-cell imaging to perform a comparative analysis of the autophagic and endocytic pathways in axons of primary cortical neurons, focusing on newly-formed organelles in the distal axon. To specifically label only newly-formed endosomes derived from the distal axon, we established a microfluidic system to compartmentally isolate neuronal axons (Fig. 1A). Neurons are plated in the proximal chamber of the microfluidic chamber and axons grow through microgrooves (900 µm in length) to reach the distal chamber (Fig. 1A). This system enabled us to add fluorescently-labeled cargos to only the distal chamber that will be endocytosed from the distal axon. To validate the compartmentalized nature of the microfluidic chamber, we immunostained for total neuron mass using neuron-specific β3-tubulin and dendrites using MAP2; total cells were labeled with the nuclear dye Hoechst. As shown in Fig. 1B, the somatodendritic domains are restricted to the proximal chamber and only axons traverse the microgrooves to reach the distal chamber. To establish a more complex environment that includes intercellular connections between neurons and astrocytes, we co-cultured neurons with astrocytes by adding primary cortical astrocytes to only the proximal chamber. After 6 DIV of co-culture, astrocytes [immunostained for astrocyte-specific glial fibrillary acidic protein (GFAP)] remained restricted to the proximal chamber and developed elaborate star-like morphologies reminiscent of structures observed in vivo (Fig. 1B). Co-culturing neurons with astrocytes in the context of the microfluidic chamber promoted a robust extension of healthy axons through the microgrooves to reach the distal chamber.

We next validated the fluidic isolation in the microfluidic chambers. By creating a difference in volume between proximal and distal chambers, a gradient of fluidic pressure is generated that restricts treatments to a particular side of the chamber. Thus, dyes added to the distal chamber can only reach the proximal chamber if transported via axons, and not by diffusion within the microgroove, and vice versa. To label newly-formed endosomes, we added BSA conjugated to Alexa Fluor 647 (abbreviated BSA-647) to either the distal chamber or the proximal chamber (Extended Data Fig. 1-1). BSA-647 is a fluid-phase cargo that is endocytosed non-specifically into endosomes and labels the spectrum of organelles along the endolysosomal pathway. We incubated neurons for 2 hr with BSA-647, washed out unincorporated BSA, and then performed live cell imaging in axons along the middle region of the microgrooves. When BSA-647 is applied to only the distal chamber, we observed robust retrograde transport of newly-formed endosomes labeled with BSA-647 in the mid-axon (Extended Data Fig. 1-1A-A’). In fact, ∼79% of BSA-647-positive puncta travelled in the retrograde direction (>1 µm displacement in 1 min), and only ∼5% of BSA-647-positive puncta travelled in the anterograde direction (>1 µm displacement in 1 min) (Extended Data Fig. 1-1A”). The remaining ∼17% of BSA-647-positive puncta did not undergo processive movement in either direction (<1 µm displacement in 1 min), and were binned as either stationary or bidirectional (Extended Data Fig. 1-1A”). These results are consistent with prior work from Hollenbeck and colleagues establishing that endocytic cargos are taken up in the distal axon and subsequently transported to the soma (Hollenbeck, 1993; Overly and Hollenbeck, 1996). By contrast, when BSA-647 is applied to only the proximal chamber, we observed a mix in the directionality of movement in the axon (Extended Data Fig. 1-1B, B’). We found that ∼41% of BSA-647-positive puncta moved in the anterograde direction, ∼26% were retrograde, and ∼33% were stationary/bidirectional (Extended Data Fig. 1-1B”). Thus, application of BSA-647 to either the distal or proximal chamber results in distinct motility patterns of endosomes in the axon, demonstrating successful fluidic isolation in the microfluidic chambers.

To measure the extent of overlap between distally-derived endosomes and autophagosomes in the axon, we cultured neurons in the microfluidic chamber that stably express GFP-LC3, a marker for autophagic organelles (Kabeya et al., 2000; Mizushima et al., 2004), and labeled newly-formed endosomes by distal addition of BSA-647 (Fig. 1C). We incubated neurons for 2 hr with distal addition of BSA-647, washed away unincorporated BSA, and performed live cell imaging in axons along the middle region of the microgrooves. Consistent with our prior work, GFP-LC3-positive autophagosomes in the axon underwent primarily retrograde movement (Fig. 1C’) (Maday et al., 2012; Maday and Holzbaur, 2014, 2016). Consistent with our validation experiment described above, BSA-647-positive endosomes also underwent processive motility primarily in the retrograde direction (Fig. 1C’). However, we observed little overlap between GFP-LC3 and endocytosed BSA-647 in the axon (Fig. 1C’). To quantify the degree of overlap, we measured the percentage of autophagosomes in axons within each microgroove that were co-positive for BSA-647. To ensure that the GFP-LC3-positive axon reached the distal chamber to endocytose BSA, we quantified only those GFP-LC3-positive axons that had at least one BSA-647-positive punctum. Strikingly, we found that only ∼1.2% of GFP-LC3-postive autophagosomes were positive for BSA-647 (Fig. 1C”). The vast majority (∼98.8%) of autophagosomes were negative for BSA-647 (Fig. 1C”). Thus, these results indicate that retrograde autophagosomes and newly-formed endosomes are remarkably distinct organelle populations in the axon.

We next wondered whether washing out the BSA -647 prior to imaging depletes an endocytic pool of BSA that would be merging with autophagosomes. To address this possibility, we performed a continuous uptake of BSA-647 in the distal chamber (Fig. 1D). For this experiment, neurons were incubated with BSA-647 in the distal chamber for 2 hr, and the BSA-647 remained present in the distal chamber during image acquisition in the mid-axon. Similar to our results with the pre-loaded BSA-647, continuous addition of BSA-647 also labeled retrograde endosomes that did not overlap with retrograde autophagosomes (Fig. 1D’). We found that 0% of GFP-LC3-positive autophagosomes per axon were co-positive for BSA-647, and 100% of autophagosomes per axon were negative for BSA-647 (Fig. 1D”). Thus, even with a continuous supply of BSA-647 endocytic cargo, we find no overlap between retrograde autophagosomes and newly endocytosed cargo in the axon.

To comprehensively examine the overlap between autophagosomes and endosomes in the axon, we tracked a population of endosomes termed signaling endosomes. Signaling endosomes are endosomes that carry active BDNF-TrkB signaling complexes from the presynaptic membrane to the cell soma to elicit changes in gene expression (Chowdary et al., 2012; Harrington and Ginty, 2013). Several recent reports have proposed that a population of signaling endosomes may merge with autophagosomes to form a signaling amphisome (Kononenko et al., 2017; Andres-Alonso et al., 2019). To explore this possibility, we incubated the distal axons with BDNF conjugated to Quantum-dots-605 for 2 hr, and then imaged in the middle of the microgrooves (Fig. 1E). As expected, the BDNF-Qdots exhibited robust retrograde transport (Fig. 1E’). However, we found little overlap between autophagosomes and BDNF-Qdots labeling signaling endosomes, and only 0.6% GFP-LC3-positive autophagosomes per microgroove were positive for BDNF-Qdots (Fig. 1E”). Thus, these data indicate that autophagosomes are predominantly not merging with signaling endosomes, and these organelle populations are distinct in axons. Combined, we find that axonal autophagosomes and newly-formed endosomes undergo retrograde transport in parallel but separate pathways.

### Retrograde autophagosomes and newly-formed endosomes exhibit distinct rates of maturation

We next investigated the maturation state of organelles in these pathways. First, we determined the extent of organelle acidification using LysoTracker, a cell-permeant dye that accumulates in acidic compartments of pH <∼6. Similar with prior reports, LysoTracker-positive organelles in the axon moved predominantly in the retrograde direction as opposed to the anterograde direction (Fig. 2A’, B’, C’, Extended Data Fig. 2-1B’”-B””) (Maday et al., 2012; Gowrishankar et al., 2017; Lie et al., 2021). This retrograde bias in motility was observed when Lysotracker was added to either the proximal (Extended Data Fig. 2-1B’”-B””) or distal chamber (Fig. 2A’, B’, C’), likely owing to its membrane-permeant chemistry. We next treated only the distal chamber with LysoTracker and quantified the percentage of autophagosomes in axons within each microgroove that were co-positive for LysoTracker (Fig. 2A). As above, we analyzed only GFP-LC3-positive axons that had at least one LysoTracker-positive puncta to ensure that the axon reached the distal chamber to access the dye. We found that nearly all (∼98%) of GFP-LC3-positive organelles were positive for LysoTracker (Fig. 2A’, A”). These results are consistent with our prior reports and indicate that autophagosomes in the axon are maturing into acidic organelles (Maday et al., 2012). By contrast, newly-formed endosomes (labeled with BSA conjugated to Alexa Fluor 488; abbreviated BSA-488) exhibited a broad spectrum in the percentage of endosomes in axons that are acidic, resulting in a mean value of ∼48% co-positivity (Fig. 2B-B”). Thus, endosomes in the axon are acidifying at uniquely distinct rates as compared with autophagosomes. Moreover, signaling endosomes labeled with BDNF-Qdots exhibited low overlap (only ∼22%) with LysoTracker (Fig. 2C-C”). This lack of acidification is consistent with their role in preserving active BDNF-TrkB signaling complexes; acidic environments attenuate TrkB signaling (Ouyang et al., 2013). Thus, retrograde autophagosomes and newly formed endosomes exhibit different degrees of acidification, providing further evidence that these pathways are composed of distinct organelle populations.

Given the high degree of heterogeneity in the acidification of newly-formed endosomes in the axon, we investigated their maturation state further by examining colocalization with markers of late endosomes and lysosomes, Rab7 and LAMP1. The switch from Rab5 to Rab7 is a critical determinant in endosomal maturation from early endosomes to late endosomes (Rink et al., 2005). LAMP1 is marker for late endosomes and lysosomes that plays a role in lysosome biogenesis and structural integrity (Eskelinen, 2006). Overlap between Rab7/LAMP1 and autophagosomes has been studied extensively in neurons (Lee et al., 2011; Maday et al., 2012; Cheng et al., 2015; Maday and Holzbaur, 2016; Kulkarni et al., 2021). In fact, nearly all LC3 compartments in the axon are positive for LAMP1 (Kulkarni et al., 2021) and Rab7 (Cheng et al., 2015). These data are consistent with the high degree of acidification (Fig. 2A’-A”), and their maturation into amphisomes. The precise maturation state of newly-formed endosomes, however, is less well understood. To investigate the degree of overlap between newly formed endosomes and markers for maturation into late endosomes, we co-transfected wild type neurons with GFP-Rab7 and LAMP1-RFP, and performed triple color live-cell imaging with newly endocytosed BSA-647 from the distal axon. Since we only counted BSA-647-positive puncta present in transfected neurons, we pooled 41 BSA-647-positive puncta counted across 22 axons. Of those BSA-647-positive puncta, ∼59% were co-positive for GFP-Rab7 (Fig. 3A-A”). This value closely reflected the ∼48% overlap with LysoTracker (Fig. 2B”), suggesting that the newly-formed endosomes that have matured into Rab7-positive late endosomes likely represent the acidified population. Surprisingly, only ∼17% of BSA-647-positive puncta were co-positive for LAMP1-RFP (5 out of 7 BSA/LAMP1-co-positive puncta were also positive for Rab7; Fig. 3A-A”). Thus, the majority of newly-formed endosomes are Rab7-positive and LAMP1-negative. Thus, while both Rab7 and LAMP1 are both markers for late endosomes, they do not overlap completely and there is a population of endosomes that is positive for only Rab7. Moreover, our data indicate distinct mechanisms of maturation between distally-formed endosomes and autophagosomes.

Paradigms established in non-neuronal cells describe extensive overlap between Rab7-positive endosomes and LAMP1 (Humphries et al., 2011). Therefore, we were surprised to find that newly-formed endosomes in the axon are positive for Rab7 and largely negative for LAMP1. We sought to further define the LAMP1-positive compartments in the axon by parsing out overlap with Rab7 based on directionality of movement. We found that total LAMP1-positive compartments have nearly equivalent movements into and out of the axon (Fig. 3B, C). On average, ∼42% of LAMP1-positive compartments in the axon exhibit anterograde motility, ∼31% exhibit retrograde motility, and the remaining ∼27% are stationary/bidirectional (Fig. 3B, C). Strikingly, Rab7 is enriched on LAMP1 compartments moving in the retrograde direction as compared with the anterograde direction (Fig. 3B, C’). We find that an average of ∼60% of retrograde LAMP1 compartments are positive for Rab7, whereas only ∼13% of anterograde LAMP1 compartments are positive for Rab7 (Fig. 3C’). In fact, the entire population of anterograde LAMP1 compartments was negative for Rab7 in ∼63% of axons (Fig. 3C”). Thus, we find different pools of LAMP1-positive organelles in the axon. We find an anterograde pool of LAMP1 compartments that are largely negative for Rab7 and may represent post-Golgi carriers described by Lie et al. that deliver lysosomal components to the distal axon (Lie et al., 2021). We also find a retrograde pool of LAMP1 compartments that are enriched for Rab7. Given the low degree of overlap between LAMP1 and newly-formed endosomes, this retrograde LAMP1-Rab7 co-positive population likely represents retrograde LC3 organelles. Common to both distally-generated autophagosomes and endosomes is the acquisition of Rab7 in distal regions of the axon.

We also analyzed these data from the Rab7 point of view. Rab7-positive puncta have roughly equivalent movements into and out of the axon, with a slight bias in the retrograde direction; ∼14% anterograde versus ∼24% retrograde (Fig. 3D). The largest population of Rab7-positive puncta were those exhibiting stationary or bidirectional movement (∼62%; Fig. 3D). We also observed an enrichment of LAMP1 on retrograde Rab7-positive compartments as compared to anterograde Rab7-positive organelles (Fig. 3D’). In fact, the entire population of retrograde Rab7 compartments was positive for LAMP1 in ∼37% of axons as compared to ∼49% of axons in which the entire anterograde population of Rab7 compartments was negative for LAMP1 (Fig. 3D”). Thus, we find a bias for Rab7-LAMP1 co-positive compartments in the axon moving in the retrograde direction as compared to the anterograde direction, suggesting a recruitment of Rab7 onto organelles in the distal axon.

### Newly endocytosed cargos are not degraded in the axon, but rather in the soma

A key question in the field of axonal biology is the extent to which lysosome-mediated degradation occurs in the axon. Thus, we determined whether newly-formed endosomes in the axon possessed proteolytic activity. For this experiment, we incubated the distal chamber for 2 hr with BSA-647 along with DQ-Green-BSA, a fluorescently labeled version of BSA that self-quenches in its native state and fluoresces with proteolytic cleavage (Fig. 4A). Consequently, DQ-Green-BSA labels proteolytically-active lysosomes. To ensure that sufficient substrate is delivered to label degradative lysosomes in the soma, we also incubated the proximal chamber with only DQ-Green-BSA (Fig. 4A). To assess the degradative state of endosomal organelles along the axon, we performed dual color live-cell imaging of BSA-647 and DQ-Green-BSA. To determine the fate of endosomes derived from the distal axon, we imaged the BSA conjugates in the proximal chamber. In the axon, we observed robust retrograde transport of endosomes positive for BSA-647, however, these organelles were negative for fluorescent DQ-Green-BSA (Fig. 4A”’). In fact, few organelles positive for DQ-Green-BSA were detected in the axon (Fig. 4A”’). The BSA-647 organelles present in the axon likely contain the DQ-Green-BSA substrate, but in an uncleaved and self-quenched form. In the soma, however, we observed a concentration of DQ-Green-BSA-positive puncta, consistent with several reports indicating the soma as the central location of proteolytically-active lysosomes in neurons (Fig. 4A’) (Gowrishankar et al., 2015; Cheng et al., 2018; Yap et al., 2018; Kulkarni et al., 2021; Lie et al., 2021). Moreover, we observed the accumulation of BSA-647 in the soma in organelles positive for DQ-Green-BSA (Fig. 4A’). Line scan analysis of BSA-647 puncta in the soma revealed extensive overlap between these two markers (Fig. 4A’, A”). Thus, BSA-647 is endocytosed in the distal axon and travels the distance of the axon to reach degradative compartments in the soma. These newly-formed endosomes in the axon largely do not yet possess proteolytic activity, and cargo do not appear to be degraded until they reach mature lysosomes in the soma.

To assess whether the presence of degradative activity in newly-formed endosomal organelles in the axon requires a longer incubation to process the DQ-BSA substrate, we performed the same experiment as above but with a 4 hr incubation with the BSA conjugates (Fig. 4B). Similar to 2 hr, retrograde organelles positive for endocytosed BSA-647 were negative for DQ-Green-BSA, and we found few DQ-Green-BSA-positive puncta in the axon (Fig. 4B”’). In the soma, however, BSA-647 and DQ-Green-BSA colocalized extensively (Fig. B’, B”). Thus, these results further support that the degradation of cargo endocytosed in the distal axon occurs predominantly in the soma.

Since several reports have indicated the presence of a population of degradative lysosomes in the axon (Farias et al., 2017; Farfel-Becker et al., 2019), we explored this possibility further and compared the distribution of DQ-BSA conjugated to either Green or Red fluorescent probes after 2 or 4 hr of incubation. As in Fig. 4, DQ-Green-BSA was enriched in the soma, and largely absent from axons at either 2 or 4 hr (Extended Data Fig. 2-1A). Similar to DQ-Green-BSA, DQ-Red-BSA was also present in the soma (Extended Data Fig. 2-1A). However, in contrast to DQ-Green-BSA, use of DQ-Red-BSA revealed a population of puncta in the axon, and the presence of these puncta was increasingly prevalent at 4 hr as compared with 2 hr (Extended Data Fig. 2-1A). These results suggested the possibility of degradative lysosomes in the axon that were revealed only with DQ-Red-BSA possibly owing to differences in signal to noise of the BSA conjugates. Thus, we reasoned that if these DQ-Red-BSA-positive puncta in the axon were in fact degradative lysosomes, these organelles should be acidic to enable robust proteolytic activity. To test this possibility, we co-added LysoTracker-Green with DQ-Red-BSA to the proximal chamber and performed dual-color imaging in the axon (Extended Data Fig. 2-1B). Interestingly, DQ-Red-BSA-positive puncta exhibited long-range processive movement in either anterograde or retrograde directions, at roughly equal percentages (Extended Data Fig. 2-1B’; 50.6% in the anterograde direction and 49.4% in the retrograde direction; n=29 microgrooves from 3 independent experiments). In contrast, ∼80% of LysoTracker-positive organelles undergoing long-range processive movement, moved in the retrograde direction (Extended Data Fig. 2-1B’, B”’, B””). Accordingly, DQ-Red-BSA puncta entering the axon in the anterograde population were largely negative for LysoTracker (Extended Data Fig. 2-1B’, B”). In fact, only ∼5% of anterograde DQ-Red-BSA-positive puncta co-labeled with LysoTracker (Extended Data Fig. 2-1B”). Moreover, the vast majority of retrograde DQ-Red-BSA-positive puncta were also negative for LysoTracker (Extended Data Fig. 2-1B”). In contrast to the axon, we observed extensive overlap between DQ-Red-BSA-positive puncta and LysoTracker in the soma (Extended Data Fig. 2-1B’). Combined, these data are inconsistent with the majority of DQ-Red-BSA puncta in the axon representing degradative lysosomes. Rather, these data suggest that DQ-Red-BSA is initially processed in a lysosome to activate fluorescence, but then after extended incubations, may be trafficked in the neuron in a non-degradative organelle derived from a lysosome (Yu et al., 2010). Our findings raise caution to the use of these dyes after long-term treatments, and underscore the need for corroborative evidence from multiple probes to validate organelle function.

### Expression of pathogenic α-synuclein increases merging between autophagosomes and newly-formed endosomes

Lastly, we wondered whether the retrograde pathways for autophagy and endocytosis remain separate in models of neurodegeneration. We hypothesized that the context of neurodegeneration might increase merging between these pathways, resulting in anomalous degradation of cargos that should be preserved. To examine this hypothesis, we expressed GFP-tagged wild type (WT) α-synuclein or a pathogenic A53T mutant form of α-synuclein, both of which are causally linked to Parkinson’s disease (PD) (Polymeropoulos et al., 1997; Singleton et al., 2003; Chartier-Harlin et al., 2004). That is, triplication of the WT α-synuclein gene or the A53T α-synuclein variant causes early onset PD in rare familial forms of the disease (Polymeropoulos et al., 1997; Singleton et al., 2003), whereas duplication of the WT α-synuclein gene causes PD with presentation similar to idiopathic PD (Chartier-Harlin et al., 2004). Alpha-synuclein is enriched in presynaptic terminals and has a proposed role in the exocytosis and endocytosis of synaptic vesicles, and intracellular trafficking (Burre et al., 2018; Huang et al., 2019). Thus, α-synuclein represents a candidate in regulating early steps of endocytosis in presynaptic terminals. For this experiment, primary cortical neurons were grown in microfluidic chambers and transfected with WT or A53T α-synuclein. To assess overlap between autophagy and endosomal pathways, neurons were co-transfected with mCherry-LC3, and BSA-647 was applied to the distal chamber; axons were imaged in the middle of the microgroove. Because mCherry is more stable in acidic organelles compared to GFP, mCherry-LC3 labels immature organelles that would be positive for GFP-LC3, and an additional population of degradative autolysosomes in which the GFP would be quenched (Kimura et al., 2007; Kulkarni et al., 2021). Given that mCherry-LC3 labels a broad spectrum of organelles, the combined population is referred to as autophagic vacuoles (AVs). We observed that expression of A53T α-synuclein increased the number of AVs present in the axon (Fig. 5A). To quantify this effect, we measured area flux defined as the number of mCherry-LC3-positive puncta detected per 100 µm axon per minute of imaging. We found that A53T α-synuclein resulted in a statistically significant increase in the area flux of AVs in the axon (Fig. 5A, B). These results are consistent with a potential role of the pathogenic form of α-synuclein impacting the autophagy pathway.

Next, we wanted to elucidate whether expression of WT or A53T α-synuclein increased the presence of newly endocytosed cargo in autophagic vacuoles. We wanted to ensure that our analysis focused on distally-derived organelles. Since mCherry-LC3 can label more mature species that may not necessarily be derived from the distal axon, we analyzed the degree of overlap from the perspective of BSA, rather than LC3. Since our prior experiments used GFP-LC3, we first generated a comparative baseline and re-analyze the experiment in Fig. 1D-D” to calculate the percentage of BSA-647-positive puncta that were positive for GFP-LC3. Consistent with our initial analysis from the LC3 perspective, the percentage of overlap from the BSA perspective was also nearly zero (Fig. 5C). Thus, at baseline, newly endocytosed cargos are not carried in GFP-LC3-positive autophagosomes. Next, we analyzed the degree of overlap between BSA-647 and mCherry-LC3, and found that only ∼4% of BSA-647-positive puncta were co-positive for mCherry-LC3 (Fig. 5D; mock). Thus, switching the labels for LC3 did not largely affect the degree of overlap between newly endocytosed cargo and autophagic vacuoles. Interestingly, expression of WT or A53T α-synuclein increased overlap between BSA-647 and mCherry-LC3, relative to the mock control (Fig. 5A, D). Expression of WT α-synuclein increased the percentage of BSA-647 puncta co-positive for mCherry-LC3 to ∼11%, and expression of A53T α-synuclein increased the percentage of BSA-647 puncta co-positive for mCherry-LC3 to ∼29% (Fig. 5D). These effects are mild given that the majority of BSA endosomes remain distinct from autophagosomes. However, these values are very striking given that our prior experiments revealed minimal overlap between these organelle populations (Fig. 1C-D”, 5C). Moreover, we find a step-wise increase in merging of these pathways in a manner corresponding to α-synuclein pathogenicity (Fig. 5D). Thus, expression of WT or A53T α-synuclein disrupts the parallel pathways for autophagy and newly-formed endosomes, raising the possibility that alterations in organelle trafficking in the axon may contribute to the progression of disease in PD and related α-synucleinopathies.

## DISCUSSION

Autophagy and endocytic pathways shuttle cargo from the distal axon to the soma largely for degradation, but the point at which they intersect in neurons is unclear. Most studies to date examine each pathway independently. Here, we systematically define the endocytic pathway relative to autophagy in axons at high temporal resolution using live cell imaging to correlate organelle dynamics with their function. We focus our study on the trafficking of newly endocytosed cargo in the distal axon. For this, we used microfluidic chambers to distally label endosomes in bulk or a sub-population of endosomes termed signaling endosomes. Using live-cell imaging in the mid-axon, we find that axonal autophagosomes and newly-formed endosomes undergo retrograde transport as distinct organelle populations (Fig. 1). We find very little overlap between autophagosomes and signaling endosomes labeled with BDNF-Qdots. We also find very little overlap between autophagosomes and a broad population of endosomes labeled with a fluid phase substrate (BSA-647) that non-specifically labels endocytic events. Moreover, autophagic and endocytic pathways exhibit different maturation states based on two criteria. First, these pathways have different degrees of acidification (Fig. 2). Autophagosomes are uniformly labeled with the acidotropic dye Lysotracker, whereas newly-formed endosomes have a broad spectrum of acidity that likely reflects the diversity of cargos (Fig. 2). Second, these pathways exhibit different degrees of association with molecular determinants of organelle maturation (Fig. 3). We and others have shown that nearly all autophagosomes are labeled with Rab7 and LAMP1 (Lee et al., 2011; Maday et al., 2012; Cheng et al., 2015; Kulkarni et al., 2021). However, newly-formed endosomes are 60% positive for Rab7 and largely negative for LAMP1 (Fig. 3). Combined, these data support that retrograde axonal autophagy and endocytic pathways are parallel but separate in neurons (Fig. 5E, model).

Our data suggest that the distal axon is a key site for the local sorting of endocytosed cargos into distinct populations of endosomes. Endocytosed BSA labels an endosomal population of heterogeneous acidification, but BDNF-Qdots are sorted into a subpopulation of endosomes that exhibit little overlap with the acidotropic dye LysoTracker (Fig. 2). In the latter case, the lack of acidification preserves signaling information as these endosomes journey from the distal axon to the soma to alter gene expression. Thus, autophagosomes that have been reported to transport BDNF-TrkB (Kononenko et al., 2017; Andres-Alonso et al., 2019) may represent only a small population of the total autophagosomes in the axon. Our results corroborate those from Jin et al. that find that cargo derived from synaptic vesicles or the plasma membrane are also sorted into different pathways for degradation in the axon terminal (Jin et al., 2018). Moreover, Jin et al. also find that these cargos derived from the distal axon remain largely separate from autophagosomes (Jin et al., 2018). Combined, these mechanisms enable neurons to distally sort *trash from treasure*. Further signals are decoded in the soma to ensure the appropriate final destination. For example, signaling endosomes carry coronin-1 which enables them to initially avoid lysosomal fate (Suo et al., 2014). Future studies will need to elucidate how the trafficking itinerary of specific endocytic cargos relative to autophagy may vary with neuronal context including developmental stage, synaptic activity, or interactions with neighboring glia.

Our findings dovetail with reports that demonstrate that the source of LAMP1 on autophagosomes may be a population of vesicles derived from the soma (Farias et al., 2017; Farfel-Becker et al., 2019; Lie et al., 2021). These anterograde LAMP1-positive vesicles are postulated to provide distal autophagosomes with lysosomal components and proteolytic enzymes, and trigger their retrograde movement to the soma (Cheng et al., 2015; Farias et al., 2017; Farfel-Becker et al., 2019; Lie et al., 2021). The precise identity of these soma-derived LAMP1-positive vesicles, however, is under investigation, and has been proposed to be either degradative lysosomes (Farias et al., 2017; Farfel-Becker et al., 2019) or Golgi-derived transport carriers that contain lysosomal components but do not yet possess proteolytic activity (Lie et al., 2021). Our work concurs with these findings in that newly formed endosomes are not a primary source of LAMP1 on autophagosomes, as newly-formed endosomes are largely deficient for LAMP1 (Fig. 3) and do not fuse with distal autophagosomes (Fig. 1).

Moreover, our observations that newly-formed endosomes are largely negative for LAMP1 are consistent with their lack of detectable degradative activity (Fig. 4). In fact, cargos endocytosed in the distal axon do not reach a degradative compartment until the soma (Fig. 4). The low levels of LAMP1 on newly-formed endosomes in the axon suggests that these organelles do not fuse with anterograde LAMP1 carriers that would provide the lysosomal components required for degradation. In contrast, axonal autophagosomes have been reported to have some degree of proteolytic activity (Farfel-Becker et al., 2019; Lie et al., 2021). These autophagosomes, however, may not be fully acidified since the GFP fluorescence of the GFP-LC3 is not quenched. Since LysoTracker can incorporate into organelles that are only mildly acidic (pH <6), proteolytic activity in maturing axonal autophagosomes may be attributed to a subset of proteases that can be active near neutral pH (Bromme et al., 1993; Dolenc et al., 1995; Yoon et al., 2021). Nonetheless, incorporation into an autophagosome likely commits cargo to a degradative fate. By contrast, the heterogeneous nature of newly-formed endosomes likely reflects distinct destinations and functions upon arrival in the soma.

Our findings also echo results from Yap et al. that defined an endosomal population in dendrites that is Rab7-positive but LAMP1-negative (Yap et al., 2018). Rab7 is required for driving dynein-based motility of endosomes to the soma for maturation and degradation (Yap et al., 2018; Yap et al., 2022). These results are very striking given that data from non-neuronal cells predict a high degree of overlap between Rab7 and LAMP1 (Humphries et al., 2011). In our study, we also observed a population of motile endosomes in the axon that are Rab7-positive and largely LAMP1-negative (Fig. 3). Thus, axons may have a shared mechanism for endosomal maturation as in dendrites. A distinguishing feature of dendrites versus axons, however, is the degree of merging between autophagic and endolysosomal pathways. In our prior study, we observed a high degree of overlap between autophagic vacuoles and endosomes in dendrites (Kulkarni et al., 2021). However, in our current study, we report very little overlap between autophagosomes and endosomes derived in the distal axon (Fig. 1). These differences may reflect compartment-specific functions that may require more localized degradative processes to regulate the synaptic proteome in dendrites as compared with axons.

We also asked whether introduction of a pathogenic protein associated with neurodegeneration can disrupt the sorting and trafficking of newly endocytosed cargos. We found that expression of WT or A53T α-synuclein led to a mild disruption in the parallel pathways and increased the presence of endocytosed cargos in retrograde autophagosomes (Fig. 5, model). Moreover, we observed a step-wise increase in the merging of these pathways in a manner corresponding to α-synuclein pathogenicity. Expression of WT α-synuclein was sufficient to increase merging as compared to the mock control, which is consistent with gene duplications/triplications of WT α-synuclein causing PD (Singleton et al., 2003; Ibanez et al., 2004). Strikingly, expression of A53T α-synuclein resulted in the largest effect (Fig. 5), which is consistent with A53T α-synuclein causing autosomal dominant, early onset PD (Polymeropoulos et al., 1997). Interestingly, this step-wise effect of expressing WT and A53T α-synuclein has been observed in other contexts (Choubey et al., 2011). Moreover, Volpicelli-Daley et al. find that endocytosis of pathogenic pre-formed fibrils of α-synuclein impairs the retrograde transport of autophagosomes and their maturation into degradative compartments (Volpicelli-Daley et al., 2014), suggesting an interaction between these pathways in models of PD.

How might the expression of WT and A53T α-synuclein lead to merging between newly formed autophagosomes and endosomes in the axon? To date, the physiological roles of α-synuclein remain poorly understood. α-Synuclein is an intrinsically disordered cytosolic protein that binds to phospholipid membranes enriched in presynaptic terminals (Burre et al., 2018; Huang et al., 2019). α-Synuclein has been proposed to play a role in the generation of curved membranes and in the exo/endocytosis of synaptic vesicles (Xu et al., 2016). Evidence also suggests a role for α-synuclein in the clustering of synaptic vesicles to regulate the releasable pool (Diao et al., 2013; Wang et al., 2014). Reports also link α-synuclein with regulating Rab GTPase homeostasis and intracellular trafficking (Dalfo et al., 2004; Gitler et al., 2008; Soper et al., 2011). Thus, through these reported functions, overexpression of WT or A53T α-synuclein may induce aberrant merging of autophagic or endosomal organelles via a dysregulation of membrane dynamics or alterations in Rab function. Consequently, endocytosed cargos that should be preserved may be getting degraded prematurely. Alternatively, BSA-labeled organelles may become active substrates for autophagy in the distal axon upon α-synuclein expression, consistent with increased levels of autophagy. To distinguish these possibilities, future studies will need to determine the exact nature of the compartments labeled with BSA. Nevertheless, alterations in organelle homeostasis and intracellular trafficking that lead to increased merging between autophagic and endosomal pathways may contribute to neurodegeneration in α-synucleinopathies.

Moreover, other proteins associated with PD are positioned at the nexus of autophagy and endocytosis in presynaptic compartments. For example, activating mutations in the leucine-rich repeat kinase 2 (LRRK2) are associated with PD (West et al., 2005). Soukup et al. find that LRRK2-mediated phosphorylation of EndophilinA switches its role from canonical functions in synaptic vesicle endocytosis to promoting autophagy in presynaptic terminals (Soukup et al., 2016). Specifically, LRRK2-mediated phosphorylation of the N-terminal BAR domain in EndophilinA promotes the formation of highly curved membranes for the assembly of autophagosome biogenesis factors (Soukup et al., 2016). Moreover, Synaptojanin 1 serves dual roles in supporting synaptic vesicle endocytosis and autophagosome formation in presynaptic terminals via differential activity of its two lipid phosphatase domains (Vanhauwaert et al., 2017); missense mutations in Synaptojanin1 are associated with PD (Krebs et al., 2013; Quadri et al., 2013). Thus, increasing evidence supports that crosstalk between autophagy and endocytosis in axon terminals may represent a point of vulnerability for neurons. Misregulation of these pathways may lead to aberrant degradation of cargos and contribute to neuronal dysfunction early in α-synucleinopathies.

## ACKNOWLEDGMENTS

This work was supported by NIH grant R01NS110716 and associated Research Supplement to Promote Diversity in Health-Related Research to SM. We thank Dr. Kelly Jordan-Sciutto and her laboratory (University of Pennsylvania, Philadelphia, PA) for providing primary rat cortical neurons. We thank members of the Maday lab for advice with experiments and preparation of the manuscript.

---

**Figure 1-1: extended data related to Figure 1.**
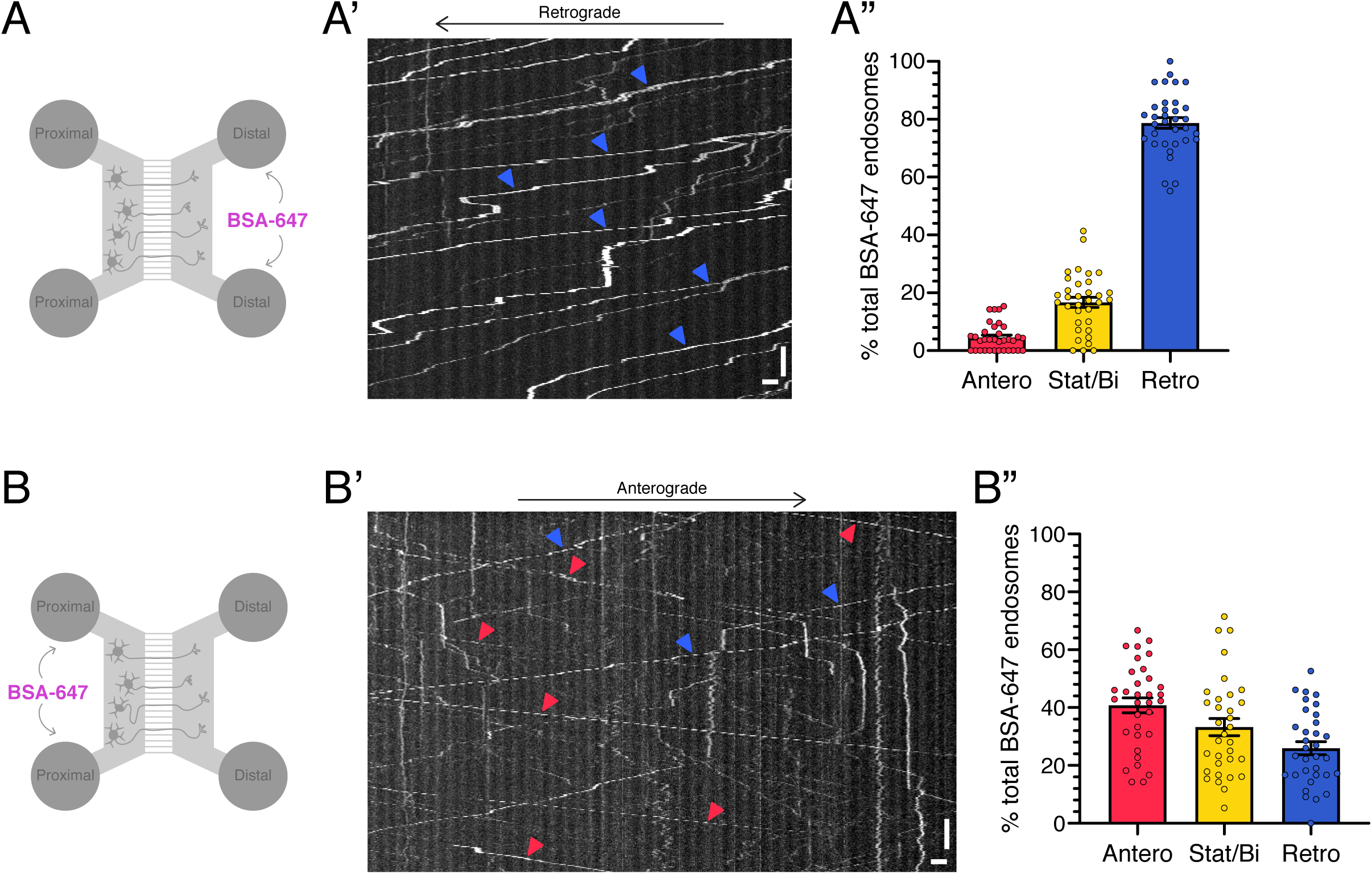
Fluidic isolation of BSA-647 in the distal chamber results in predominantly retrograde motility, but fluidic isolation of BSA-647 in the proximal chamber results in long-range motility in both anterograde and retrograde directions. **(A-A”)** Live cell imaging of newly endocytosed BSA-647 in axons of monocultured primary rat cortical neurons. BSA-647 was added to only the distal chamber. **(A)** Schematic of experimental setup. **(A’)** Kymograph analysis of newly endocytosed BSA-647 motility in the axon. Vertical bar, 15 sec. Horizontal bar, 3 µm. **(A”)** Corresponding quantitation of the directionality of BSA-647-positive puncta in the axon (means ± SEM; n=34 microgrooves from 3 independent experiments; 9-14 DIV). **(B-B”)** Live cell imaging of newly endocytosed BSA-647 in axons of primary cortical neurons. BSA-647 was added to only the proximal chamber. **(B)** Schematic of experimental setup. **(B’)** Kymograph analysis of newly endocytosed BSA-647 motility in the axon. Vertical bar, 15 sec. Horizontal bar, 3 µm. **(B”)** Corresponding quantitation of the directionality of BSA-647-positive puncta in the axon (means ± SEM; n=33 microgrooves from 3 independent experiments; 8-14 DIV).

**Figure 2-1: extended data related to Figure 2 and Figure 4.**
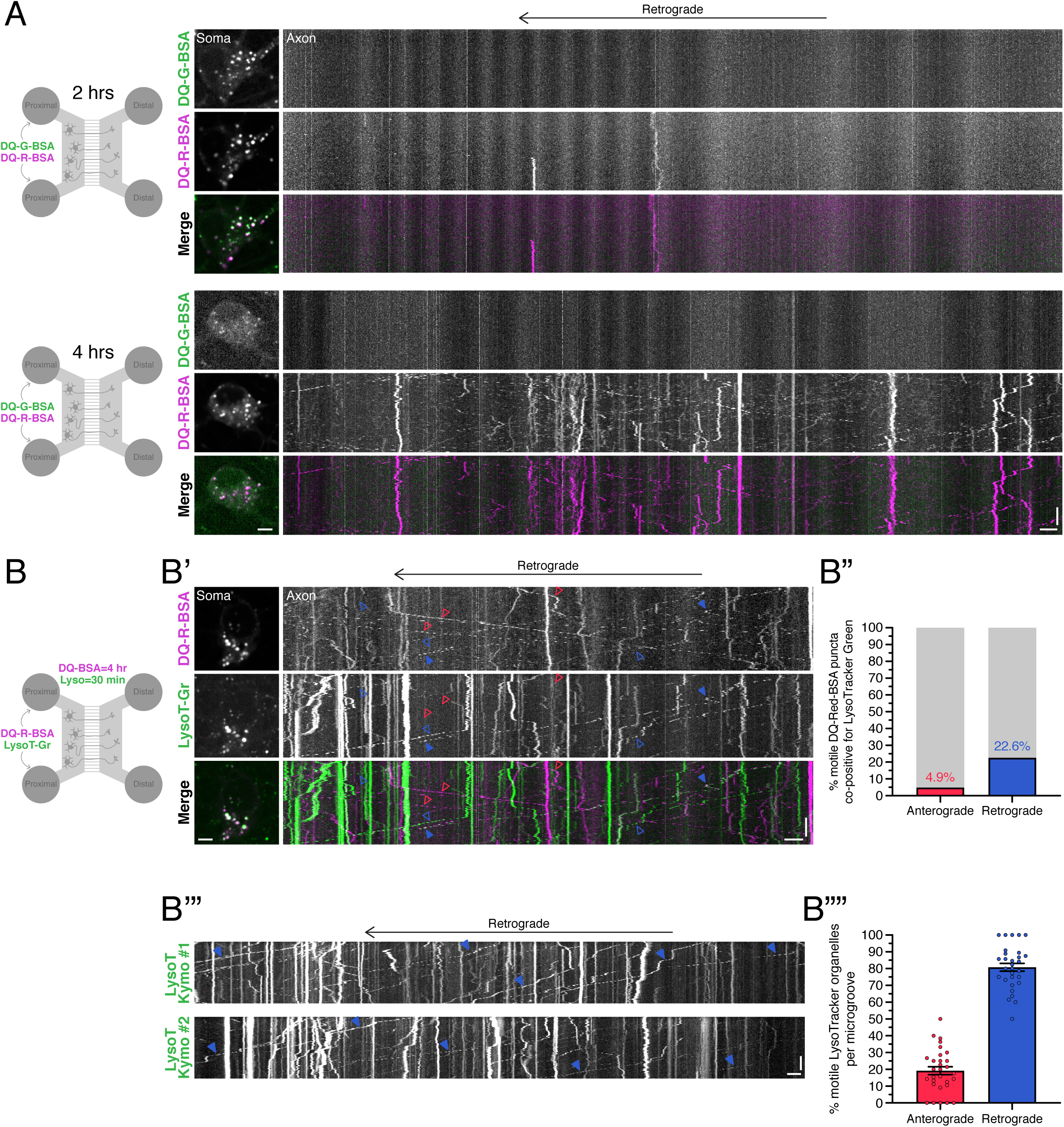
Longer incubations with DQ-Red-BSA label compartments that are not acidic in nature. **(A)** Live cell imaging of compartments labeled with DQ-Green-BSA and DQ-Red-BSA in primary cortical neurons. BSA conjugates are added to only the proximal chamber for 2 hr or 4 hr. Shown are the experimental setup, *en face* images of the soma, and kymographs of the axon. Horizontal bars, 5 µm. Vertical bar, 30 seconds. **(B-B””)** Live cell imaging of compartments labeled with DQ-Red-BSA and LysoTracker-Green in axons of primary cortical neurons. DQ-Red-BSA is added to only the proximal chamber for 4 hr; Lysotracker-Green is added during the final 30 minutes of incubation to only the proximal chamber. **(B)** Schematic of the experimental setup. (B’) *En face* images of the soma and kymographs of DQ-Red-BSA and Lysotracker-Green motility in the axon of primary cortical neurons. Red arrowheads denote anterograde puncta; open red arrowheads denote puncta positive for DQ-Red-BSA and negative for Lysotracker-Green. Blue arrowheads denote retrograde puncta; closed blue arrowheads denote puncta co-positive for DQ-Red-BSA and Lysotracker-Green and open blue arrowheads denote puncta positive for only Lysotracker-Green and not DQ-Red-BSA. Horizontal bars, 5 µm. Vertical bar, 30 seconds. **(B”)** Corresponding quantification of the percentage of motile DQ-Red-BSA puncta that are positive for LysoTracker based on directionality (percentage based on a pool of 163 anterograde DQ-Red-BSA-positive puncta or 159 retrograde DQ-Red-BSA-positive puncta counted from 29 microgrooves from 3 independent experiments; 8-11 DIV). **(B’”)** Kymographs of LysoTracker-Green motility in two separate axons of primary cortical neurons. Blue arrowheads denote retrograde puncta positive for LysoTracker-Green. Horizontal bar, 5 µm. Vertical bar, 30 seconds. **(B’’’’)** Corresponding quantification of the directionality of LysoTracker-positive puncta undergoing long-range motility (means ± SEM; n=30 microgrooves from 3 independent experiments; 8-11 DIV). Percentages per microgroove are determined from an average of ∼12 LysoTracker-positive puncta.

## REFERENCES

1. Andres-Alonso M et al. (2019) SIPA1L2 controls trafficking and local signaling of TrkB-containing amphisomes at presynaptic terminals. Nat Commun 10:5448.

2. Bromme D, Bonneau PR, Lachance P, Wiederanders B, Kirschke H, Peters C, Thomas DY, Storer AC, Vernet T (1993) Functional expression of human cathepsin S in Saccharomyces cerevisiae. Purification and characterization of the recombinant enzyme. J Biol Chem 268:4832–4838.

3. Burre J, Sharma M, Sudhof TC (2018) Cell Biology and Pathophysiology of alpha-Synuclein. Cold Spring Harb Perspect Med 8.

4. Chartier-Harlin MC, Kachergus J, Roumier C, Mouroux V, Douay X, Lincoln S, Levecque C, Larvor L, Andrieux J, Hulihan M, Waucquier N, Defebvre L, Amouyel P, Farrer M, Destee A (2004) Alpha-synuclein locus duplication as a cause of familial Parkinson’s disease. Lancet 364:1167–1169.

5. Cheng XT, Zhou B, Lin MY, Cai Q, Sheng ZH (2015) Axonal autophagosomes recruit dynein for retrograde transport through fusion with late endosomes. J Cell Biol 209:377–386.

6. Cheng XT, Xie YX, Zhou B, Huang N, Farfel-Becker T, Sheng ZH (2018) Characterization of LAMP1-labeled nondegradative lysosomal and endocytic compartments in neurons. J Cell Biol 217:3127–3139.

7. Choubey V, Safiulina D, Vaarmann A, Cagalinec M, Wareski P, Kuum M, Zharkovsky A, Kaasik A (2011) Mutant A53T alpha-synuclein induces neuronal death by increasing mitochondrial autophagy. J Biol Chem 286:10814–10824.

8. Chowdary PD, Che DL, Cui B (2012) Neurotrophin signaling via long-distance axonal transport. Annu Rev Phys Chem 63:571–594.

9. Dalfo E, Gomez-Isla T, Rosa JL, Nieto Bodelon M, Cuadrado Tejedor M, Barrachina M, Ambrosio S, Ferrer I (2004) Abnormal alpha-synuclein interactions with Rab proteins in alpha-synuclein A30P transgenic mice. J Neuropathol Exp Neurol 63:302–313.

10. Diao J, Burre J, Vivona S, Cipriano DJ, Sharma M, Kyoung M, Sudhof TC, Brunger AT (2013) Native alpha-synuclein induces clustering of synaptic-vesicle mimics via binding to phospholipids and synaptobrevin-2/VAMP2. Elife 2:e00592.

11. Dolenc I, Turk B, Pungercic G, Ritonja A, Turk V (1995) Oligomeric structure and substrate induced inhibition of human cathepsin C. J Biol Chem 270:21626–21631.

12. Dong A, Kulkarni VV, Maday S (2019) Methods for Imaging Autophagosome Dynamics in Primary Neurons. Methods Mol Biol 1880:243–256.

13. Eskelinen EL (2006) Roles of LAMP-1 and LAMP-2 in lysosome biogenesis and autophagy. Mol Aspects Med 27:495–502.

14. Farfel-Becker T, Roney JC, Cheng XT, Li S, Cuddy SR, Sheng ZH (2019) Neuronal Soma-Derived Degradative Lysosomes Are Continuously Delivered to Distal Axons to Maintain Local Degradation Capacity. Cell Rep 28:51–64 e54.

15. Farias GG, Guardia CM, De Pace R, Britt DJ, Bonifacino JS (2017) BORC/kinesin-1 ensemble drives polarized transport of lysosomes into the axon. Proc Natl Acad Sci U S A 114:E2955–E2964.

16. Gitler AD, Bevis BJ, Shorter J, Strathearn KE, Hamamichi S, Su LJ, Caldwell KA, Caldwell GA, Rochet JC, McCaffery JM, Barlowe C, Lindquist S (2008) The Parkinson’s disease protein alpha-synuclein disrupts cellular Rab homeostasis. Proc Natl Acad Sci U S A 105:145–150.

17. Gowrishankar S, Wu Y, Ferguson SM (2017) Impaired JIP3-dependent axonal lysosome transport promotes amyloid plaque pathology. J Cell Biol 216:3291–3305.

18. Gowrishankar S, Yuan P, Wu Y, Schrag M, Paradise S, Grutzendler J, De Camilli P, Ferguson SM (2015) Massive accumulation of luminal protease-deficient axonal lysosomes at Alzheimer’s disease amyloid plaques. Proc Natl Acad Sci U S A 112:E3699–3708.

19. Hangen E, Cordelieres FP, Petersen JD, Choquet D, Coussen F (2018) Neuronal Activity and Intracellular Calcium Levels Regulate Intracellular Transport of Newly Synthesized AMPAR. Cell Rep 24:1001–1012 e1003.

20. Harrington AW, Ginty DD (2013) Long-distance retrograde neurotrophic factor signalling in neurons. Nat Rev Neurosci 14:177–187.

21. Hill SE, Kauffman KJ, Krout M, Richmond JE, Melia TJ, Colon-Ramos DA (2019) Maturation and Clearance of Autophagosomes in Neurons Depends on a Specific Cysteine Protease Isoform, ATG-4.2. Dev Cell 49:251-266 e258.

22. Hollenbeck PJ (1993) Products of endocytosis and autophagy are retrieved from axons by regulated retrograde organelle transport. J Cell Biol 121:305–315.

23. Huang M, Wang B, Li X, Fu C, Wang C, Kang X (2019) alpha-Synuclein: A Multifunctional Player in Exocytosis, Endocytosis, and Vesicle Recycling. Front Neurosci 13:28.

24. Humphries WHt, Szymanski CJ, Payne CK (2011) Endo-lysosomal vesicles positive for Rab7 and LAMP1 are terminal vesicles for the transport of dextran. PLoS One 6:e26626.

25. Ibanez P, Bonnet AM, Debarges B, Lohmann E, Tison F, Pollak P, Agid Y, Durr A, Brice A (2004) Causal relation between alpha-synuclein gene duplication and familial Parkinson’s disease. Lancet 364:1169–1171.

26. Jin EJ, Kiral FR, Ozel MN, Burchardt LS, Osterland M, Epstein D, Wolfenberg H, Prohaska S, Hiesinger PR (2018) Live Observation of Two Parallel Membrane Degradation Pathways at Axon Terminals. Curr Biol 28:1027–1038 e1024.

27. Kabeya Y, Mizushima N, Ueno T, Yamamoto A, Kirisako T, Noda T, Kominami E, Ohsumi Y, Yoshimori T (2000) LC3, a mammalian homologue of yeast Apg8p, is localized in autophagosome membranes after processing. EMBO J 19:5720–5728.

28. Kimura S, Noda T, Yoshimori T (2007) Dissection of the autophagosome maturation process by a novel reporter protein, tandem fluorescent-tagged LC3. Autophagy 3:452–460.

29. Kononenko NL, Classen GA, Kuijpers M, Puchkov D, Maritzen T, Tempes A, Malik AR, Skalecka A, Bera S, Jaworski J, Haucke V (2017) Retrograde transport of TrkB-containing autophagosomes via the adaptor AP-2 mediates neuronal complexity and prevents neurodegeneration. Nat Commun 8:14819.

30. Krebs CE, Karkheiran S, Powell JC, Cao M, Makarov V, Darvish H, Di Paolo G, Walker RH, Shahidi GA, Buxbaum JD, De Camilli P, Yue Z, Paisan-Ruiz C (2013) The Sac1 domain of SYNJ1 identified mutated in a family with early-onset progressive Parkinsonism with generalized seizures. Hum Mutat 34:1200–1207.

31. Kulkarni VV, Anand A, Herr JB, Miranda C, Vogel MC, Maday S (2021) Synaptic activity controls autophagic vacuole motility and function in dendrites. J Cell Biol 220.

32. Lee S, Sato Y, Nixon RA (2011) Lysosomal proteolysis inhibition selectively disrupts axonal transport of degradative organelles and causes an Alzheimer’s-like axonal dystrophy. J Neurosci 31:7817–7830.

33. Lie PPY, Yang DS, Stavrides P, Goulbourne CN, Zheng P, Mohan PS, Cataldo AM, Nixon RA (2021) Post-Golgi carriers, not lysosomes, confer lysosomal properties to pre-degradative organelles in normal and dystrophic axons. Cell Rep 35:109034.

34. Maday S, Holzbaur EL (2014) Autophagosome biogenesis in primary neurons follows an ordered and spatially regulated pathway. Dev Cell 30:71–85.

35. Maday S, Holzbaur EL (2016) Compartment-Specific Regulation of Autophagy in Primary Neurons. J Neurosci 36:5933–5945.

36. Maday S, Wallace KE, Holzbaur EL (2012) Autophagosomes initiate distally and mature during transport toward the cell soma in primary neurons. J Cell Biol 196:407–417.

37. Malik BR, Maddison DC, Smith GA, Peters OM (2019) Autophagic and endo-lysosomal dysfunction in neurodegenerative disease. Mol Brain 12:100.

38. Mizushima N, Yamamoto A, Matsui M, Yoshimori T, Ohsumi Y (2004) In vivo analysis of autophagy in response to nutrient starvation using transgenic mice expressing a fluorescent autophagosome marker. Mol Biol Cell 15:1101–1111.

39. Ouyang Q, Lizarraga SB, Schmidt M, Yang U, Gong J, Ellisor D, Kauer JA, Morrow EM (2013) Christianson syndrome protein NHE6 modulates TrkB endosomal signaling required for neuronal circuit development. Neuron 80:97–112.

40. Overly CC, Hollenbeck PJ (1996) Dynamic organization of endocytic pathways in axons of cultured sympathetic neurons. J Neurosci 16:6056–6064.

41. Polymeropoulos MH, Lavedan C, Leroy E, Ide SE, Dehejia A, Dutra A, Pike B, Root H, Rubenstein J, Boyer R, Stenroos ES, Chandrasekharappa S, Athanassiadou A, Papapetropoulos T, Johnson WG, Lazzarini AM, Duvoisin RC, Di Iorio G, Golbe LI, Nussbaum RL (1997) Mutation in the alpha-synuclein gene identified in families with Parkinson’s disease. Science 276:2045–2047.

42. Quadri M et al. (2013) Mutation in the SYNJ1 gene associated with autosomal recessive, early-onset Parkinsonism. Hum Mutat 34:1208–1215.

43. Rink J, Ghigo E, Kalaidzidis Y, Zerial M (2005) Rab conversion as a mechanism of progression from early to late endosomes. Cell 122:735–749.

44. Sidibe DK, Vogel MC, Maday S (2022) Organization of the autophagy pathway in neurons. Curr Opin Neurobiol 75:102554.

45. Singleton AB et al. (2003) alpha-Synuclein locus triplication causes Parkinson’s disease. Science 302:841.

46. Soper JH, Kehm V, Burd CG, Bankaitis VA, Lee VM (2011) Aggregation of alpha-synuclein in S. cerevisiae is associated with defects in endosomal trafficking and phospholipid biosynthesis. J Mol Neurosci 43:391–405.

47. Soukup SF, Kuenen S, Vanhauwaert R, Manetsberger J, Hernandez-Diaz S, Swerts J, Schoovaerts N, Vilain S, Gounko NV, Vints K, Geens A, De Strooper B, Verstreken P (2016) A LRRK2-Dependent EndophilinA Phosphoswitch Is Critical for Macroautophagy at Presynaptic Terminals. Neuron 92:829–844.

48. Stavoe AK, Hill SE, Hall DH, Colon-Ramos DA (2016) KIF1A/UNC-104 Transports ATG-9 to Regulate Neurodevelopment and Autophagy at Synapses. Dev Cell 38:171–185.

49. Suo D, Park J, Harrington AW, Zweifel LS, Mihalas S, Deppmann CD (2014) Coronin-1 is a neurotrophin endosomal effector that is required for developmental competition for survival. Nat Neurosci 17:36–45.

50. Vanhauwaert R et al. (2017) The SAC1 domain in synaptojanin is required for autophagosome maturation at presynaptic terminals. EMBO J 36:1392–1411.

51. Volpicelli-Daley LA, Gamble KL, Schultheiss CE, Riddle DM, West AB, Lee VM (2014) Formation of alpha-synuclein Lewy neurite-like aggregates in axons impedes the transport of distinct endosomes. Mol Biol Cell 25:4010–4023.

52. Wang L, Das U, Scott DA, Tang Y, McLean PJ, Roy S (2014) alpha-synuclein multimers cluster synaptic vesicles and attenuate recycling. Curr Biol 24:2319–2326.

53. West AB, Moore DJ, Biskup S, Bugayenko A, Smith WW, Ross CA, Dawson VL, Dawson TM (2005) Parkinson’s disease-associated mutations in leucine-rich repeat kinase 2 augment kinase activity. Proc Natl Acad Sci U S A 102:16842–16847.

54. Winckler B, Faundez V, Maday S, Cai Q, Guimas Almeida C, Zhang H (2018) The Endolysosomal System and Proteostasis: From Development to Degeneration. J Neurosci 38:9364–9374.

55. Xu J, Wu XS, Sheng J, Zhang Z, Yue HY, Sun L, Sgobio C, Lin X, Peng S, Jin Y, Gan L, Cai H, Wu LG (2016) alpha-Synuclein Mutation Inhibits Endocytosis at Mammalian Central Nerve Terminals. J Neurosci 36:4408–4414.

56. Yap CC, Digilio L, McMahon LP, Garcia ADR, Winckler B (2018) Degradation of dendritic cargos requires Rab7-dependent transport to somatic lysosomes. J Cell Biol 217:3141–3159.

57. Yap CC, Digilio L, McMahon LP, Wang T, Winckler B (2022) Dynein Is Required for Rab7-Dependent Endosome Maturation, Retrograde Dendritic Transport, and Degradation. J Neurosci 42:4415–4434.

58. Yoon MC, Solania A, Jiang Z, Christy MP, Podvin S, Mosier C, Lietz CB, Ito G, Gerwick WH, Wolan DW, Hook G, O’Donoghue AJ, Hook V (2021) Selective Neutral pH Inhibitor of Cathepsin B Designed Based on Cleavage Preferences at Cytosolic and Lysosomal pH Conditions. ACS Chem Biol 16:1628–1643.

59. Yu L, McPhee CK, Zheng L, Mardones GA, Rong Y, Peng J, Mi N, Zhao Y, Liu Z, Wan F, Hailey DW, Oorschot V, Klumperman J, Baehrecke EH, Lenardo MJ (2010) Termination of autophagy and reformation of lysosomes regulated by mTOR. Nature 465:942–946.

